# Highly accurate metagenome-assembled genomes from human gut microbiota using long-read assembly, binning, and consolidation methods

**DOI:** 10.1101/2024.05.10.593587

**Authors:** Daniel M. Portik, Xiaowen Feng, Gaetan Benoit, Daniel J. Nasko, Benjamin Auch, Samuel J. Bryson, Raul Cano, Martha Carlin, Anabelle Damerum, Brett Farthing, Jonas R. Grove, Moutusee Islam, Kyle W. Langford, Ivan Liachko, Kristopher Locken, Hayley Mangelson, Shuiquan Tang, Siyuan Zhang, Christopher Quince, Jeremy E. Wilkinson

**Affiliations:** PacBio, 1305 O’Brien Dr, Menlo Park, CA 94025 USA; Department of Data Sciences, Dana-Farber Cancer Institute, Boston, MA 02215 USA; Department of Biomedical Informatics, Harvard Medical School, Boston, MA 02115 USA; Sequence Bioinformatics, Department of Computational Biology, Institut Pasteur, Paris, France; Phase Genomics, 1617 8^th^ Ave, Seattle, WA 98109 USA; The BioCollective LLC., 5650 Washington St., Denver, CO 80216 USA; Zymo Research Corporation, 17062 Murphy Ave, Irvine, CA 92614 USA; Organisms and Ecosystems, Earlham Institute, Norwich, UK; Gut Microbes and Health, Quadram Institute, Norwich, UK; School of Biological Sciences, University of East Anglia, Norwich, UK; Warwick Medical School, University of Warwick, Coventry, UK

## Abstract

Long-read metagenomic sequencing is a powerful approach for cataloging the microbial diversity present in complex microbiomes, including the human gut microbiome. We performed a deep-sequencing experiment using PacBio HiFi reads to obtain metagenome-assembled genomes (MAGs) from a pooled human gut microbiome. We performed long-read metagenome assembly using two methods (hifiasm-meta, metMDBG), used improved bioinformatic and proximity ligation binning strategies to cluster contigs and identify MAGs, and developed a novel framework to compare and consolidate MAGs (pb-MAG-mirror). We found proximity ligation binning yielded more MAGs than bioinformatic binning, but our novel comparison framework resulted in higher MAG yields than either binning strategy individually. In total, from 255 Gbp of total HiFi data we produced 595 total MAGs (including 175 high-quality MAGs) using hifiasm-meta, and 547 total MAGs (including 277 high-quality MAGs) with metaMDBG. Hifiasm-meta assembled almost twice as many strain-level MAGs as metaMDBG (246 vs. 156), but both assembly methods produced up to five strains for a species. Approximately 85% of the MAGs were assigned to known species, but we recovered >35 high-quality MAGs that represent uncultured diversity. Based on strict similarity scores, we found 125 MAGs were unequivocally shared across the assembly methods at the strain level, representing ∼22% of the total MAGs recovered per method. Finally, we detected more total viral sequences in the metaMDBG assembly versus the hifiasm-meta assembly (∼6,700 vs. ∼4,500). Overall, we find the use of HiFi sequencing, improved metagenome assembly methods, and complementary binning strategies is highly effective for rapidly cataloging microbial genomes in complex microbiomes.

## INTRODUCTION

The human gut microbiome contains a diversity of microbes that potentially impact health and disease (Lynch & Pederson 2016; Wang & Jia 2016; Duvallet et al. 2017; Gilbert et al. 2018). Despite considerable efforts to catalog the microbial diversity present in the human gut, large numbers of species and strains remain uncultured and undetected (Lagier et al. 2018; Nayfach et al. 2019; Almeida et al. 2019; Forster et al. 2019; Pasolli et al. 2019; Zou et al. 2019; Almeida et al. 2021). The pace of biodiversity discovery is limited using traditional isolation and culturing methods, but it can be greatly accelerated using metagenomic sequencing. Metagenome assembly is a powerful approach for reconstructing the genomic contents of species contained in a microbiome sample (Tyson et al. 2004). Historically, the contigs produced by metagenome assemblies often represent small fragments of microbial genomes, and binning methods are used to group contigs into putative genomes. The genomes obtained through assembly and binning are commonly referred to as metagenome-assembled genomes (MAGs). Metagenome assembly based on short-read sequencing generally requires substantial effort to produce high-quality MAGs (Bowers et al. 2017; Chen et al. 2020). For example, repetitive genomic regions and interspecies genomic overlaps are difficult to resolve using short reads, and this generally leads to highly fragmented assemblies (Arumugam et al. 2019). Furthermore, binning can introduce major errors by grouping contigs belonging to different species or strains into MAGs (Chen et al. 2020). This can result in chimeric MAGs or even MAGs with human contamination, which can have severe negative effects on downstream analyses (see Gihawi et al. 2023).

Long-read sequencing can overcome many of the challenges associated with metagenome assembly (Kolmogorov et al. 2020; Feng et al. 2022; Albertsen 2023; Benoit et al. 2023; Agustinho et al. 2024). The most popular long-read sequencing platforms are those produced by Pacific Biosciences (PacBio) and Oxford Nanopore Technologies (ONT). PacBio HiFi sequencing produces highly accurate consensus reads (>Q20, median Q30) that are 10–20 kb in length (Wenger et al. 2019). Several studies have demonstrated that HiFi sequencing generally produces more total MAGs and higher quality MAGs than short-read sequencing (Priest et al. 2021; Gehrig et al. 2022; Meslier et al. 2022; Eisenhofer et al. 2023; Orellana et al. 2023; Tao et al. 2023; Zhang et al. 2023), and that HiFi sequencing outperforms ONT for metagenome assembly (Meslier et al. 2022; Sereika et al. 2022). The high accuracy of HiFi reads has led to the development of two new metagenome assembly methods, hifiasm-meta (Feng et al. 2022) and metaMDBG (Benoit et al. 2023), which perform better than previous methods such as HiCanu (Nurk et al. 2020) and metaFlye (Kolmogorov et al. 2020). Hifiasm-meta phases reads based on single nucleotide variants and constructs the string graph using only intra-haplotype read overlaps. Heuristics to keep contained reads with ambiguous phasing and to resolve tangles using either unitig read coverages or topology are implemented in graph cleaning steps. By contrast, metaMDBG uses de-Bruijn graph assembly in a mimizer-space and a progressive abundance-based filtering strategy to simplify strain complexity. Performing metagenome assembly with these HiFi-specific methods routinely produces complete, circular MAGs, and sometimes in large numbers (Bickhart et al. 2022; Feng et al. 2022; Kato et al. 2022; Kim et al. 2022; Zhang et al. 2022; Benoit et al. 2023; Jiang et al. 2023; Saak et al. 2023; Schaerer et al. 2023; Masuda et al. 2024). For example, using hifiasm-meta Kim et al. (2022) recovered 102 complete, circular MAGs from five human gut microbiome samples (including 39 MAGs that remain uncultured), and a reanalysis of data from Bickhart et al. (2022) with metaMDBG recovered 447 high-quality MAGs from a single sheep gut microbiome, including 266 circular MAGs (Benoit et al. 2023).

Although metagenome assembly with HiFi sequencing data can produce complete MAGs, additional genomes will be represented by two or more contigs. Reconstructing these additional MAGs requires the use of binning algorithms. A majority of bioinformatic binning methods perform clustering using tetranucleotide frequencies, depth of coverage, or deep learning methods (Sczyrba et al. 2017; Meyer et al. 2022), and most were designed for short-read assemblies. When applied to long-read assemblies, these methods can produce unexpected or suboptimal results. To address this, new long-read binning algorithms have been developed, including MetaBCC-LR (Wickramarachchi et al. 2020), LRBinner (Wickramarachchi & Lin 2022), GraphMB (Lamurias et al. 2022), and most recently SemiBin2 (Pan et al. 2023).

Metagenomic assembly workflows often include multiple bioinformatic binning methods, and the results from multiple binning algorithms are typically merged. This key step de-replicates the bin sets from alternative methods, resulting in improvements to MAG quality and yield (Olm et al. 2017; Uritskiy et al. 2018). In addition to bioinformatic approaches, proximity ligation information (e.g., Hi-C) can also be used to bin contigs (Burton et al. 2014; Press et al. 2017; DeMaere & Darling 2019). The ProxiMeta platform (Press et al. 2017), MetaCC (Du & Sun 2023), and metaBAT-LR (Ho et al. 2023) can perform binning using Hi-C data and a set of assembled contigs. Proximity ligation also allows mobile elements to be associated with their respective host genomes, offering a unique advantage. Proximity ligation binning methods have been successfully applied to long-read metagenome assemblies, presumably improving MAG yields relative to bioinformatic binning (Bickhart et al. 2022; Saak et al. 2023). However, no studies have performed a systematic comparison of the MAG sets produced from bioinformatic and proximity ligation binning using long reads.

Here, we performed a deep-sequencing experiment on a pooled human gut microbiome to produce a catalog of highly resolved MAGs. This pooled gut sample is particularly challenging for metagenome assembly, as the number of species and strains is higher than what is typically found in a gut microbiome. We obtained HiFi data from the PacBio Sequel IIe and Revio systems. We assembled the combined sequencing dataset using hifiasm-meta and metaMDBG, and performed binning using bioinformatic and proximity ligation approaches. For bioinformatic binning, we created a new version of a workflow designed to process long-read metagenome assemblies (HiFi-MAG-Pipeline), and we used the ProxiMeta platform to perform proximity ligation binning and association of mobile elements. We also developed a new algorithm (pb- MAG-mirror) which compares the contents of MAGs obtained from two alternate binning approaches (based on the same set of starting contigs), and consolidates the MAGs into a single non-redundant set. In an effort to characterize tradeoffs in performance, we compared the number, quality, and taxonomy of MAGs obtained from the various combinations of assembly and binning methods. Finally, we performed a downsampling experiment to understand the effects of total data on MAG recovery. Overall, our study illustrates how HiFi sequencing can be used to obtain highly complete MAGs from the human gut microbiome, accelerating our ability to catalog the microbial diversity in this complex system.

## RESULTS

### Sequencing datasets

Our combined PacBio HiFi sequencing dataset consists of 34.7 million HiFi reads and 255.9 Gb of total data, with a mean QV of 42.5 (Table 1). We obtained 17.1 million HiFi reads (120.8 Gb) from six 8M SMRT Cells on the Sequel IIe system, and 17.5 HiFi reads (135.0 Gb) from two 25M SMRT Cells on the Revio system. The Sequel IIe HiFi reads display a mean length of 7.0 kb and mean QV of 40.5 (Supplemental Fig. S1); yields across cells ranged from 2.30–3.33 million reads and 14.1–27.8 Gb total data. The Revio HiFi reads had a similar mean length of 7.7 kb but a higher mean QV of 45.8. Importantly, predicted read QV scores were higher in the Revio dataset despite a fewer mean number of passes (18 versus 24; Supplemental Fig. S1). Both Revio runs were highly consistent in yielding 8.7–8.8 million HiFi reads and 67.0–67.9 Gb total data.

**Table 1.** Summary of HiFi sequencing datasets.

| Dataset | Sequencing level | HiFi reads (million) | HiFi yield (Gb) | Mean read length | Mean QV |
| --- | --- | --- | --- | --- | --- |
| Revio | 2 SMRT Cells (25M) | 17.56 | 135.06 | 7,690 | 45.8 |
| Sequel IIe | 6 SMRT Cells (8M) | 17.15 | 120.85 | 7,046 | 40.2 |
| Combined | - | 34.71 | 255.91 | 7,724 | 42.5 |

### Metagenome assembly and binning

We performed metagenome assembly using two HiFi-specific methods: hifiasm-meta and metaMDBG (Fig. 1). The assembly with hifiasm-meta resulted in 34,117 contigs with an N50 of 458 kbp and total assembly size of 2.76 Gbp. We found that metaMDBG produced 66,952 contigs with an N50 of 306 kbp and total assembly size of 2.75 Gbp. The runtime performance differed between the assembly methods. Using hifiasm-meta took 5 days and 5.8 hours wall time using 64 threads (7767 CPU hrs) with peak memory of 713 GB, whereas metaMDBG required 21.13 hours wall time (1352 CPU hrs) and 12.7 GB peak memory (using 64 threads).

**Figure 1.**
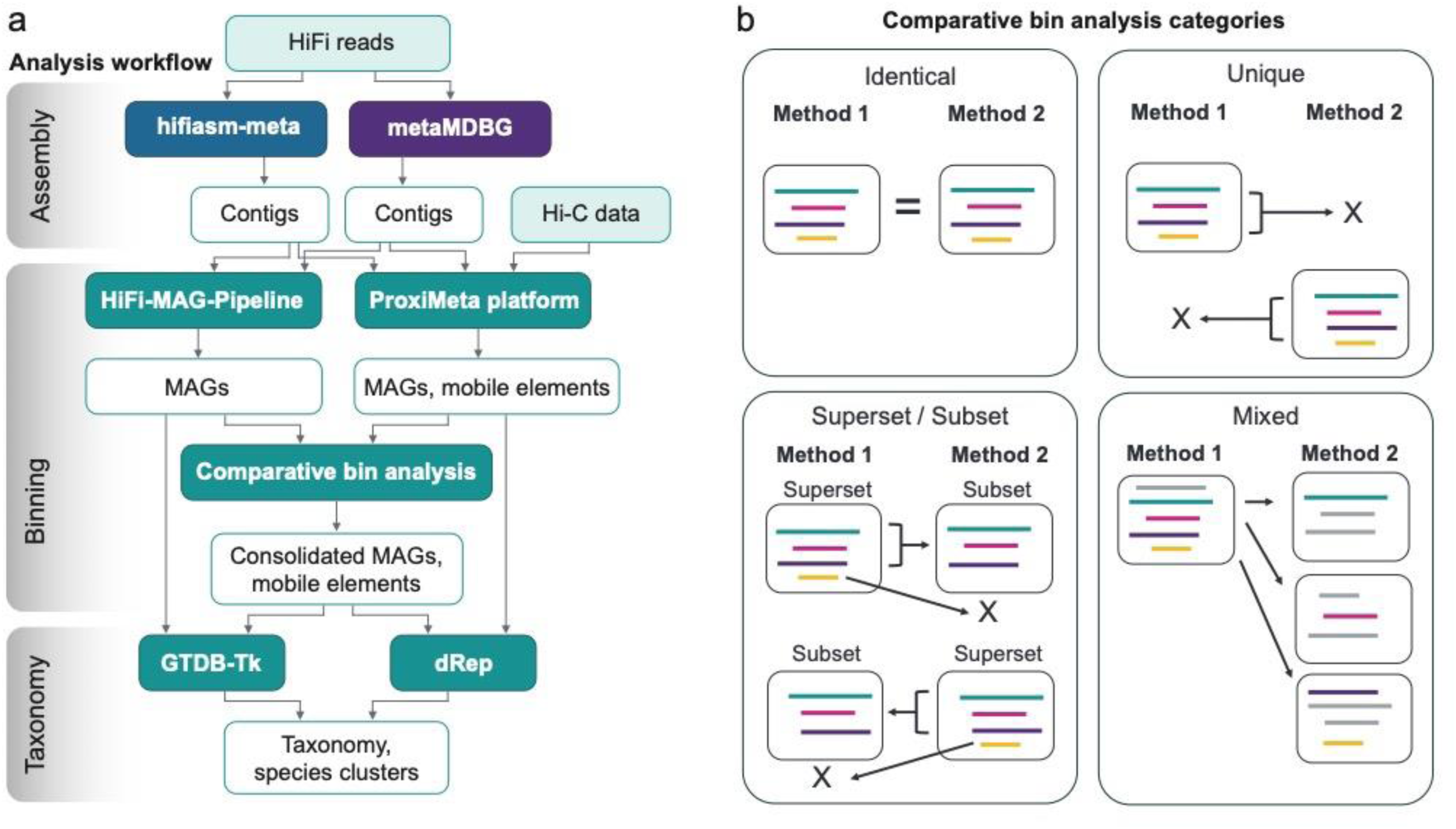
**(a)** Visual depiction of analysis steps involved with metagenome assembly, binning, and taxonomy. **(b)** Illustration of the categories used to compare MAGs with pb-MAG-mirror. Within a category, each box represents an individual MAG and the colored lines represent the contigs contained in the MAG. A full description of each category is provided in the Methods.

To obtain MAGs from the assembled contigs, we performed bioinformatic binning using the HiFi-MAG-Pipeline and proximity ligation binning using ProxiMeta (Fig. 1). We categorized MAGs as medium-quality or high-quality (MQ-MAG, HQ-MAG, respectively) based on standards proposed by the Genomic Standards Consortium (Bowers et al. 2017). The HQ-MAGs require ≥90% completeness (based on universal single-copy genes - SCGs) and ≤5% contamination, whereas MQ-MAGs fall below these criteria but display ≥50% SCG completeness and ≤10% contamination. Overall, we obtained large numbers of total MAGs and HQ-MAGs across all the assembly and binning combinations. Specifically, using bioinformatic binning we obtained 485 and 480 total MAGs for hifiasm-meta and metaMDBG, respectively (Fig. 2, Table 2). Comparing the binning strategies, we found ProxiMeta produced more total MAGs than bioinformatic binning, with 522 and 492 total MAGs for hifiasm-meta and metaMDBG, respectively (Fig. 2, Table 2). This represents an 8% and 3% increase in total MAGs for the two assembly methods. Across both binning strategies, the total number of MAGs produced by hifiasm-meta is higher than metaMDBG. However, we found metaMDBG has a higher proportion of HQ-MAGs relative to hifiasm-meta across both binning strategies (bioinformatic: 239 vs. 157; proximity ligation: 258 vs. 166; Fig 2). Both assembly methods recovered similar numbers of SC-HQ-MAGs, but the numbers differed across binning strategy (Table 2). The first step of the HiFi-MAG-Pipeline was designed to identify highly complete, single-contig MAGs using a “completeness-aware” strategy. Based on this approach, we recovered 98 and 96 SC-HQ-MAGs from hifiasm-meta and metaMDBG, respectively, with 70 and 74 being circular. The ProxiMeta workflow does not include a parallel initial step, and as a result we recovered 79 and 70 SC-HQ-MAGs (from hifiasm-meta and metaMDBG, respectively, both with 59 circular). For ProxiMeta we observed several cases in which one or more small contigs (<50kb) were binned with complete bacterial chromosomes (identified as SC-HQ-MAGs using HiFi-MAG-Pipeline), and they were subsequently excluded from being counted as SC-HQ-MAGs. The smaller contigs may possibly represent unannotated mobile elements. Overall, ProxiMeta produced higher HQ and MQ-MAG yields than the HiFi-MAG-Pipeline.

**Figure 2.**
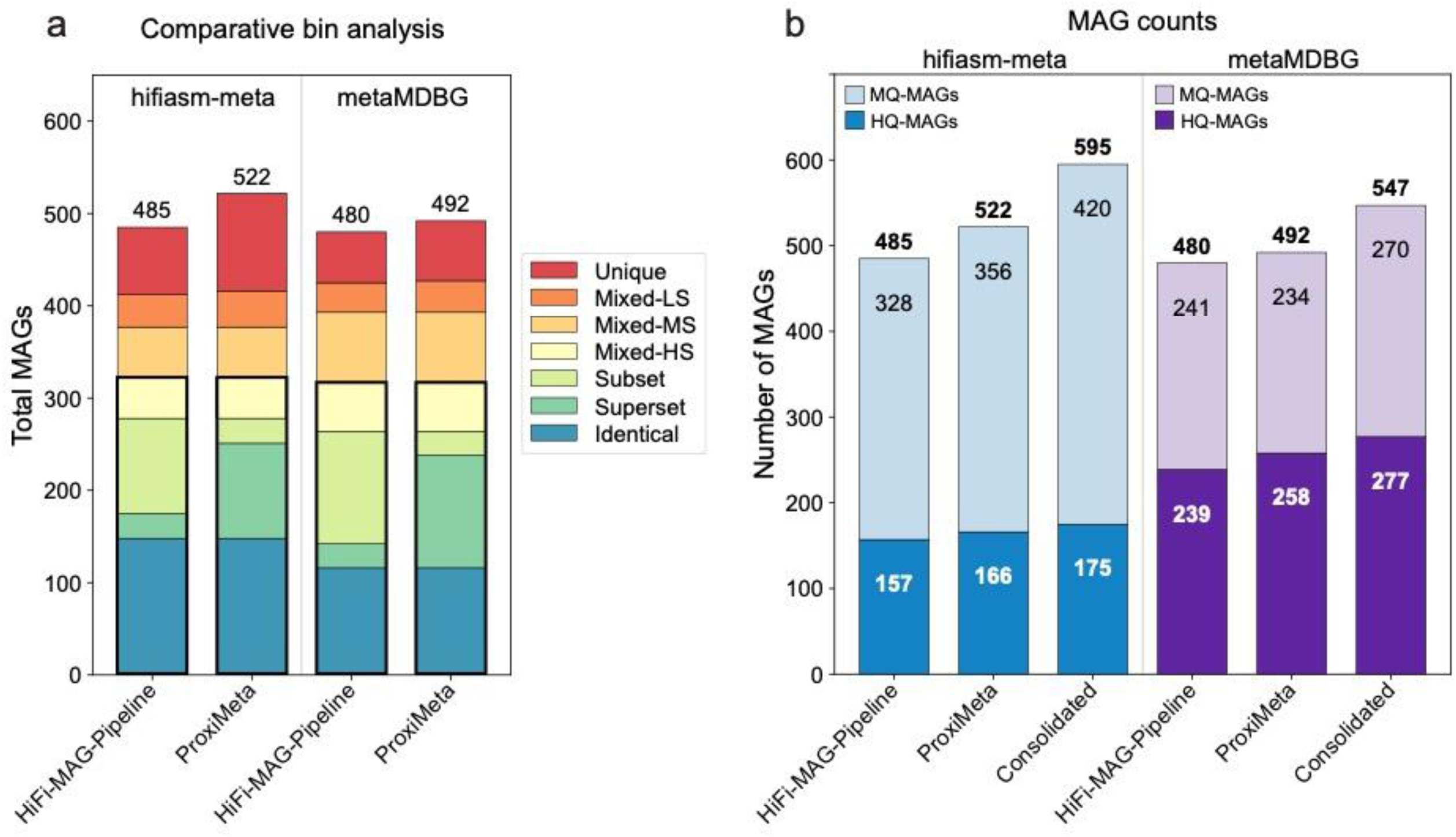
**(a)** Assignment of MAGs to categories from pb-MAG-mirror across the two assembly methods. Stacked barplot colors represent the number of MAGs occurring in different categories; total MAG numbers are shown on top. Bold outline highlights categories that are considered identical or nearly identical across methods. (**b)** Number of MAGs recovered for each assembly and binning method combination. HQ-MAGs require ≥90% single-copy genes (SCG) completeness and ≤5% contamination, whereas MQ-MAGs fall below the HQ thresholds but display ≥50% SCG completeness and ≤10% contamination. Total MAG numbers are shown above, with colors showing MAG counts exclusive to each category.

**Table 2.** Counts of MAGs by quality for each dataset and binning method combination.

| Assembly method | Binning method | SC-HQ-MAGs | HQ-MAGs | MQ-MAGs | Total MAGs |
| --- | --- | --- | --- | --- | --- |
| hifiasm-meta | HiFi-MAG-Pipeline | 70/98 | 157 | 328 | 485 |
|  | ProxiMeta | 59/79 | 166 | 356 | 522 |
|  | Consolidated | 61/83 | 175 | 420 | 595 |
| metaMDBG | HiFi-MAG-Pipeline | 74/96 | 239 | 241 | 480 |
|  | ProxiMeta | 59/70 | 258 | 234 | 492 |
|  | Consolidated | 61/73 | 277 | 270 | 547 |

The HQ-MAG category requires >90% completeness and <5% contamination, whereas MQ-MAGs fall below those criteria but display >50% completeness and <10% contamination. Total MAGs = HQ + MQ-MAGs. SC-HQ-MAGs are a subcategory of HQ-MAGs with one contig present. For SC-HQ-MAGs, the first number represents genomes that are circular, whereas the second number represents all genomes (linear or circular).

We compared and consolidated the bioinformatic and proximity ligation bin sets using our new method, pb-MAG-mirror. Our analysis with pb-MAG-mirror resulted in more MQ- and HQ-MAGs than either binning approach individually (Fig. 2). The analysis assigned MAGs to four comparison categories, including identical, superset/subset, unique, and high-, medium- and low-similarity mixed (Fig. 1). Three of these categories involved identifying highly similar pairs of MAGs across the two binning strategies (e.g., the identical, superset/subset, and high-similarity mixed categories; Fig. 1), whereas the unique, medium- and low-similarity mixed categories identified distinct or poorly overlapping MAGs. These classifications allowed us to identify the proportion of highly similar MAGs shared across binning methods, which guided the consolidation step (see Methods). Our analysis with pb-MAG-mirror produced 595 and 547 total consolidated MAGs for hifiasm-meta and metaMDBG, representing 11–22% more total MAGs than bioinformatic or proximity ligation binning individually. We found that 61–66% of total MAGs were classified as highly similar across the bioinformatic and proximity ligation binning methods (Fig. 2, Supplementary Table S1). This result indicates that a majority of MAGs contain highly similar contents across the binning approaches. By contrast, we found approximately 18– 23% of the total MAGs were assigned to the medium and low-similarity mixed categories, which had little to no overlap in MAG composition. These MAGs tended to have lower completeness scores and higher contamination scores relative to other categories (Supplemental Fig. S2).

Finally, the number of unique MAGs varied across binning and assembly methods, ranging from 11–20% of the total MAGs. ProxiMeta produced more unique MAGs than HiFi-MAG-Pipeline, and the highest number of unique MAGs occurred for the hifiasm-meta and ProxiMeta combination (Fig. 2, Supplementary Table S1). We investigated the characteristics of the unique MAGs from each binning method, and found the unique MAGs from HiFi-MAG-Pipeline and ProxiMeta had similar completeness scores (∼70%, for both assembly methods; Supplementary Fig. S2). However, the unique MAGs from HiFi-MAG-Pipeline displayed a higher average number of contigs (hifiasm-meta: 17 vs. 9 contigs; metaMDBG: 18 vs. 13 contigs) and a lower average depth of coverage (hifiasm-meta: 21x vs. 64x; metaMDBG: 27x vs. 87x), relative to ProxiMeta (Supplementary Fig. S2). In the consolidated MAG set, a major contribution comes from the inclusion of the unique MAGs from both methods. The combined unique MAGs represent 30% and 22% of the total consolidated MAGs for hifiasm-meta and metaMDBG, respectively.

### Effects of total data

To investigate the effects of total data on MAG yield, we downsampled our Sequel IIe dataset to lower data levels (Supplementary Table S2). Overall, we found the patterns from the full dataset were recapitulated in the downsampled datasets. For example, across both assembly methods we found the consolidated MAG set resulted in the greatest numbers of MQ- and HQ-MAGs, followed by proximity ligation binning, then bioinformatic binning (Supplementary Figs. 3, 4; Supplementary Table S3). In the full sequencing datasets, hifiasm-meta produced more total MAGs than metaMDBG (Fig. 2), but this pattern was not consistent across the downsampled datasets. We found metaMDBG produced more total MAGs in 14% of the downsampled comparisons, most often in conjunction with the HiFi-MAG-Pipeline (Supplementary Figs. S3, S4, Table S3). However, across all data levels we found metaMDBG produced more HQ-MAGs, and fewer MQ-MAGs, than hifiasm-meta (Fig. 3). We note our smallest downsampled dataset contained only ∼360,000 HiFi reads and 3Gb total data, yet it resulted in up to 73 total MAGs and 10 HQ-MAGs (Supplementary Table S3). Looking across all data levels (including our full dataset), we found a predictable logarithmic relationship between total data and the number of HQ or MQ-MAGs in the consolidated MAG sets (Fig. 3; Supplementary Table S4). These trendlines indicate that although additional sequencing could further increase MAG yields, there are diminishing returns relative to lower data levels.

**Figure 3.**
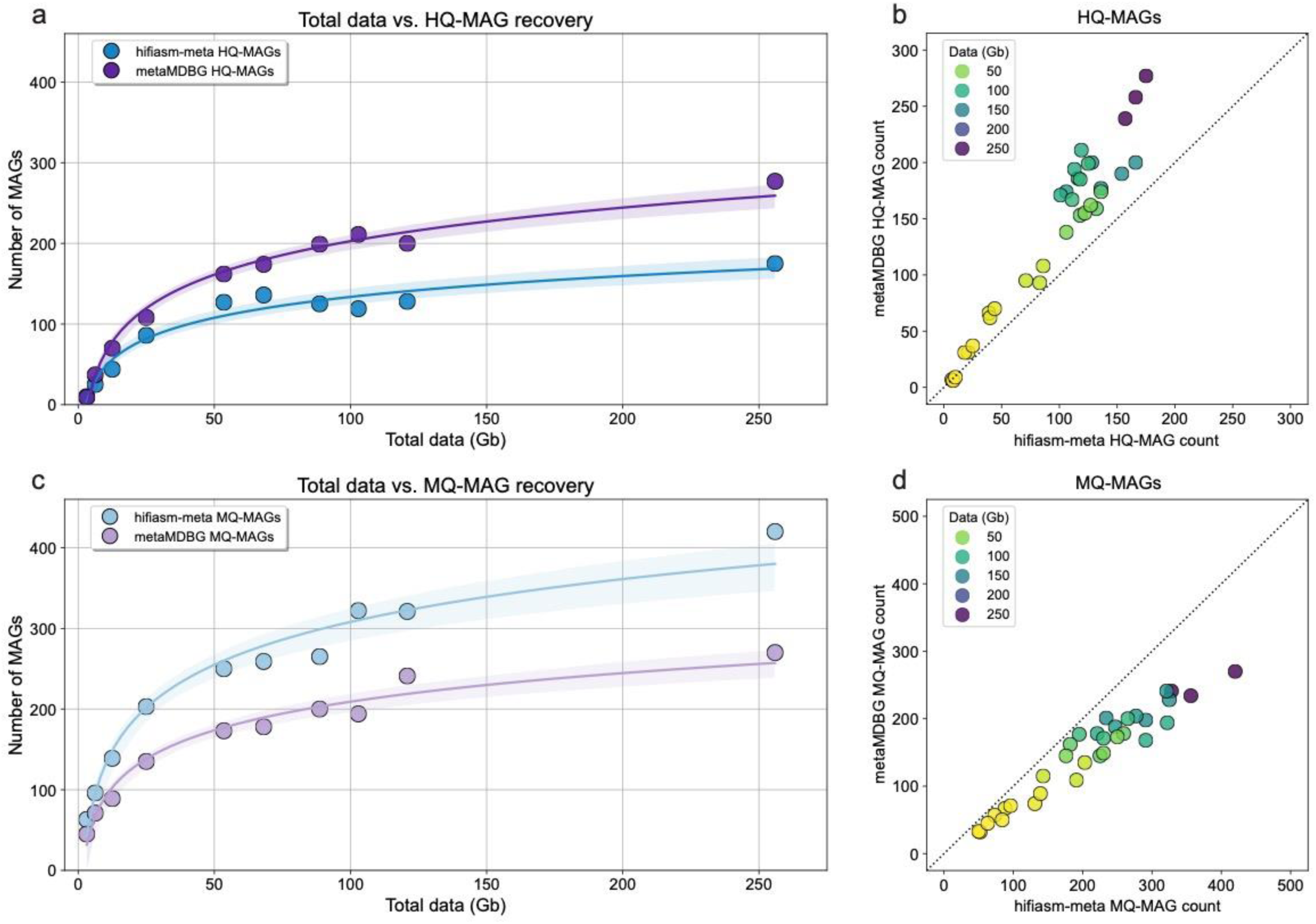
**(a)** Number of HQ-MAGs for the consolidated MAG set across downsampled datasets. (**b)** Number of HQ-MAGs obtained from hifiasm-meta versus metaMDBG across all binning methods and downsampled datasets. (**c)** Number of MQ-MAGs for the consolidated MAG set across downsampled datasets. (**d)** Number of MQ-MAGs obtained from hifiasm-meta versus metaMDBG across all binning methods and downsampled datasets.

### Taxonomic diversity and strain-level variation

The number of species assigned by GTDB-Tk ranged from 313–385 species across assembly and binning combinations, with the highest number of species occurring in the consolidated MAG sets for hifiasm-meta (n=364) and metaMDBG (n=385; Table 3). The number of MAGs that were not assigned to the species level ranged from 65–96 across the method combinations, representing 13–17% of the total MAGs. The highest numbers again occurred in the consolidated MAG sets, with 96 MAGs not assigned to the species rank for hifiasm-meta (16%) and 75 for metaMDBG (14%). Given the high number of unassigned MAGs, the species counts from GTDB-Tk represent a conservative estimate of the total species diversity. The dRep results support this notion, as we recovered more species-level clusters across assembly and binning combinations, ranging from 385–455 (Table 3). The highest numbers of dRep species clusters were also found for the consolidated MAG sets, with 449 clusters for hifiasm-meta and 455 for metaMDBG (Fig. 4).

**Figure 4.**
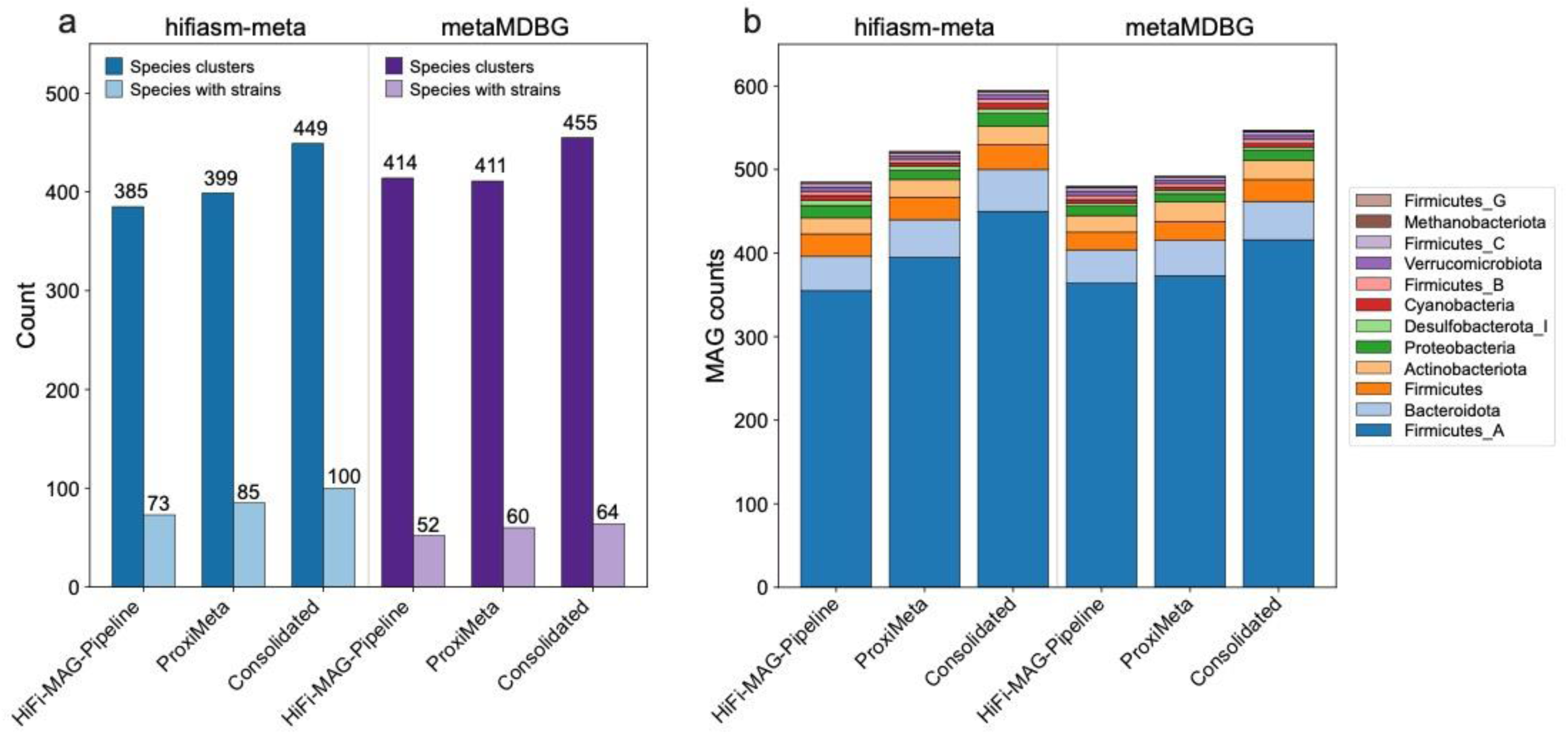
**(a)** Total number of species and species containing multiple strains, for each assembly and binning combination. Counts are based on dRep clustering results. (**b)** Aggregated taxonomic counts of MAGs assigned at the phylum rank.

**Table 3.** Estimates of species and strain information obtained by reference-based taxonomic assignment (GTDB-Tk) and MAG clustering (dRep).

| Assembly method | Binning method | GTDB-Tk |  |  | dRep |  |  |
| --- | --- | --- | --- | --- | --- | --- | --- |
|  |  | Total species | Species with strains | Total strain-level MAGs | Unique clusters | Clusters with strains | Total strain-level MAGs |
| hifiasm-meta | HiFi-MAG-Pipeline | 313 | 67 | 158 | 385 | 73 | 173 |
|  | ProxiMeta | 329 | 82 | 196 | 399 | 85 | 208 |
|  | Consolidated | 364 | 97 | 233 | 449 | 100 | 246 |
| metaMDBG | HiFi-MAG-Pipeline | 347 | 50 | 115 | 414 | 52 | 118 |
|  | ProxiMeta | 351 | 60 | 137 | 411 | 60 | 141 |
|  | Consolidated | 385 | 64 | 152 | 455 | 64 | 156 |

We observed a large proportion of strain-level variation in our MAG sets, particularly for method combinations including hifiasm-meta. Based on dRep species clusters, we found 73–100 species represented by multiple strains across the binning methods for hifiasm-meta, with only 52–64 species displaying multiple strains for metaMDBG (Fig. 4; Table 3). For hifiasm-meta, we found a total of 246 strain-level MAGs in the consolidated MAG set, representing 40% of the total MAGs (Table 3). A large proportion of these strain-level genomes for hifiasm-meta were classified as MQ-MAGs (68%). By contrast, we only found 156 strain-level MAGs for metaDBG, representing 28% of the total MAGs (Table 3). Of these, 52% were classified as MQ-MAGs. In the consolidated MAG sets there were many species containing two strains (hifiasm-meta: 68; metaMDBG: 46), several with three strains (hifiasm-meta: 20; metaMDBG: 10), but only a few displaying four (hifiasm-meta: 10; metaMDBG: 6) or five strains (hifiasm-meta: 2; metaMDBG: 2). Across all species with 2–5 strains, approximately ∼65% contained at least one HQ-MAG, ∼25% contained two or more HQ-MAGs, ∼15% contained three HQ-MAGs, and none contained four or more HQ-MAGs (Supplementary Table S5). These results indicate that assembling multiple HQ-MAGs at the strain level is currently possible, but that strain-level variation is typically captured as one HQ-MAG plus one or more incomplete genomes. Among the species clusters displaying four to five strains, there are 12 for hifiasm-meta and 8 for metaMDBG (Supplementary Table S6). There are 15 high-strain-diversity species in the combined set: *Adlercreutzia celatus_A/equolifaciens, Agathobacter faecis, Agathobaculum butyriciproducens, Bacteroides uniformis, Blautia_A massiliensis, CAG-41 sp900066215, Copromonas sp000435795, Dorea_A longicatena, Dysosmobacter sp001916835, Faecalibacterium prausnitzii_D, Faecalibacterium prausnitzii_G, Faecalibacterium sp900539945, Lachnospira eligens_A/sp003451515, Phocaeicola dorei/vulgatus,* and *Ruminococcus_B gnavus*. We note that in addition to these high-strain-diversity species, there are an additional 26 species which contain three strains each.

Our main phylogenetic and taxonomic analyses were focused on the consolidated MAG sets for hifiasm-meta and metaMDBG (Fig. 4). At the phylum level, the MAG assignments were dominated by Firmicutes_A (hifiasm-meta: 450; metaMDBG: 416), followed by Bacteroidota and Firmicutes (hifiasm-meta: 50 and 30, respectively; metaMDBG: 46 and 26). Within Firmicutes_A, representation was highest among the families Lachnospiraceae (hifiasm-meta: 184; metaMDBG: 153), Oscillospiraceae (hifiasm-meta: 61; metaMDBG: 68), Ruminococcaceae (hifiasm-meta: 54; metaMDBG: 57), and Acutalibacteraceae (hifiasm-meta: 33; metaMDBG: 31), but included 33 families in total. Within Bacteroidota seven families were represented, with Bacteroidaceae (hifiasm-meta: 24; metaMDBG: 19) and Rikenellaceae (hifiasm-meta: 13; metaMDBG: 12) displaying the greatest number of MAGs. The phyla with the least amount of representation included Methanobacteria, Cyanobacteria, Desulfobacterota_I, Firmicutes_B, Firmicutes_C, Firmicutes_G, and Verrucomicrobiota (hifiasm-meta: 2, 6, 6, 5, 4, 0, and 5 MAGs, respectively; metaMDBG: 1, 4, 4, 5, 4, 1, and 5 MAGs). Based on GTDB-Tk, we determined 96 of the consolidated MAGs were not assigned to the species level for hifiasm-meta, and 75 were not assigned for metaMDBG. Of these, 21 from hifiasm-meta and 16 from metaMDBG were not assignable to the genus level. Of the set of MAGs not assigned at the species level, we found 16 and 36 were categorized as HQ-MAGs for hifiasm-meta and metaMDBG, respectively (Supplementary Table S7). A majority of these novel HQ-MAGs occur in the phylum Firmicutes_A (hifiasm-meta: 10; metaMDBG: 28), and within this phylum most occur in the orders Lachnospirales and Oscillospirales (Supplementary Table S7). In total, the novel HQ-MAGs recovered from hifiasm-meta and metaMDBG represent 33 distinct genera distributed across thirteen orders and six phyla.

### Comparison of assembly methods

We performed comparisons of hifiasm-meta and metaMDBG at the contig and MAG-level, using several approaches. We performed an alignment of the contigs across assemblers, and found 69% of bases were shared at the 98% identity threshold, and 63% of bases were shared at the 99% identity threshold. We also compared large contigs (>500kb) using Mash and FracMinHash to understand contig similarity and containment (Supplementary Table S8). At a 99% mash similarity threshold (e.g., strain-level), we found 13.8% and 16.3% of large contigs were matched across hifiasm-meta and metaMDBG, with 13.2% and 14.4% being contained. At 95% for mash and 80% for FracMinHash, the number of contig matches for hifiasm-meta and metaMDBG was 39% and 53%, and 44% and 33% were contained. Finally. for 90% for mash and 70% for FracMinHash, the number of matched long contigs for hifiasm-meta and metaMDBG was 51% and 64%, with 47% and 33% contained.

We mapped the HiFi reads to the contigs and to the consolidated MAGs for both assembly methods, and found similar numbers of reliably mapped reads in both cases (Supplementary Table S9). Approximately 78% of HiFi reads were reliably mapped to the contigs (for both assembly methods), and 61% and 62% were mapped to the consolidated MAGs (hifiasm-meta and metaMDBG, respectively). These results indicate a large proportion of the information contained in the HiFi reads was represented in the contigs, with a somewhat smaller proportion represented in the final MAGs.

To understand how much species diversity was assembled into genomes, we searched for 16S genes in the HiFi reads and compared them to the 16S genes contained within MAGs. We found 108,830 reads contained full length 16S rRNA genes, and greedy clustering yielded 3,772 species-level OTUs (with a minimum of two reads each). We found 1,645 OTUs had more than 10 reads of support. Of the 1,645 OTUs with high coverage, 436 (26.5%) were also found in the consolidated HQ-MAG set (Fig. 5). When considering the MQ- and HQ-MAGs, we found a total of 778 OTUs represented (47.3%).

**Figure 5.**
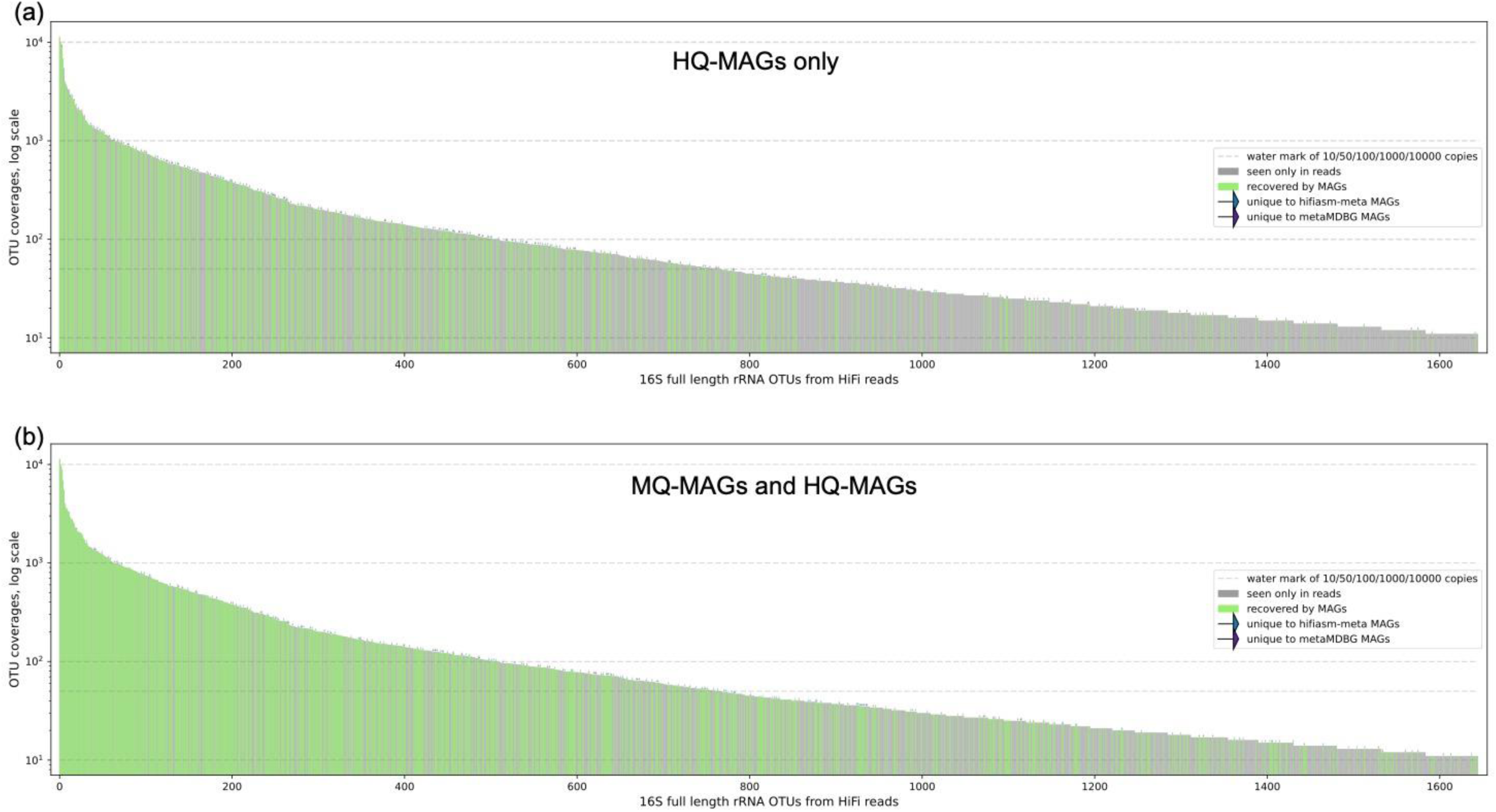
Barplots showing the 16S OTUs identified from the HiFi reads, based on 99% identity clustering. Relative abundance is represented by the height of the bars (log coverage). OTUs only occurring in the HiFi reads are shown in grey, whereas green indicates the OTU was also detected in a MAG. The MAG sets consist of the consolidated MAGs from both hifiasm-meta and metaMDBG, with results shown for **(a)** HQ-MAGs only or (**b)** all MQ- and HQ-MAGs.

We estimated the number of MAGs that were unequivocally shared across the assembly methods, based on the consolidated MAG sets. We used a combination of aligned sequence lengths and ANI to measure MAG similarity, and explored the effects of different values on the number of matches (Supplemental Fig. S5). Ultimately, we defined an unequivocal match as requiring ≥90% of the total bases per MAG to be aligned with ≥99% ANI (Fig. 6), which is representative of a strain-level match. Based on these criteria, we found a total of 125 MAGs that were unequivocally shared across the assembly methods, representing 21–23% of the total MAGs for each method (Fig. 6, Supplementary Table S10). These unequivocally shared MAGs had higher average percent completeness scores (97% vs. 73–84%), lower average numbers of contigs (3–7 vs. 9–12), and higher average depths of coverages (186–192x vs. 72–78x) relative to the unmatched MAGs from each assembly method (Fig. 6). We determined 115 of the shared MAGs (92%) were classified as HQ-MAGs, but we also found matched MAGs with as low as 71% completeness.

**Figure 6.**
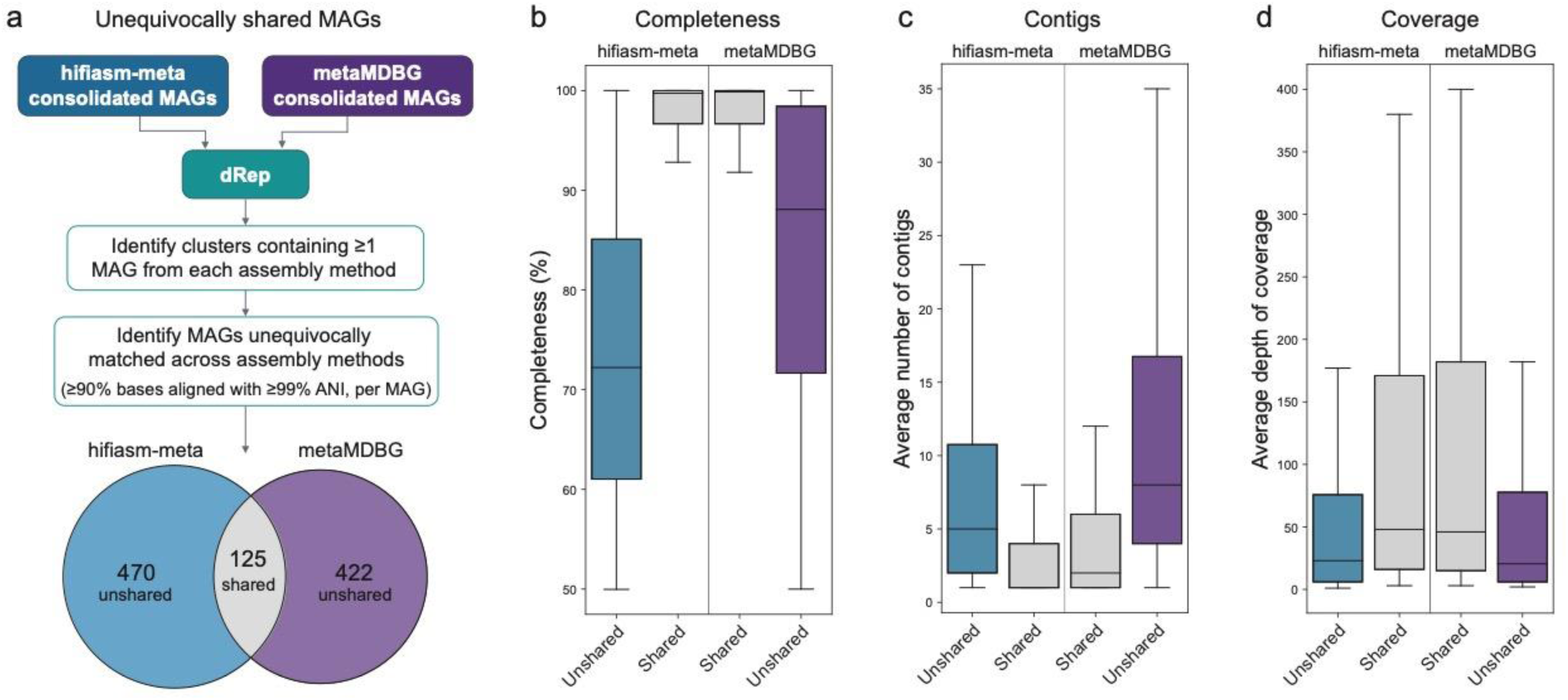
**(a)** Visual overview of workflow used to identify MAGs unequivocally shared across the assembly methods. Barplots summarizing (**b)** percent completeness, (**c)** average number of contigs per MAG, and (**b)** depth of coverage for MAGs in the shared (grey) and not shared categories (blue: hifiasm-meta, purple: metaMDBG). Outliers not shown.

### Mobile element association and viral detection

We used ProxiMeta, geNomad, and VirSorter2 to detect viral, proviral, and plasmid sequences in the contigs from both assembly methods. For both assemblers geNomad detected the most total viral contigs (hifiasm-meta 4,551, metaMDBG 6,679), followed closely by VirSorter2 (hifiasm-meta 4,264, metaMDBG 6,203), and then ProxiMeta (hifiasm-meta 2,175, metaMDBG 2,501; Supplemental Table S11). For a given assembly, a large proportion of viral and proviral contigs were annotated by both VirSorter2 and geNomad (62–80%; Supplementary Fig. S6). In addition to viral and proviral categories, ProxiMeta and geNomad annotated plasmid sequences. We found geNomad recovered more plasmids for metaMDBG (11,983) relative to hifiasm-meta (6,827). ProxiMeta identified fewer total plasmid sequences than geNomad but differentiated between plasmids and integrated plasmids. For metaMDBG ProxiMeta detected 129 plasmids and 1,878 integrated plasmids, whereas for hifiasm-meta it found 164 plasmids and 2,268 integrated plasmids (Supplementary Table S11). In addition to identifying mobile element sequences, ProxiMeta’s ProxiPhage algorithm predicts if a given mobile sequence is interacting with a microbial bin as a host. We found that ProxiMeta identified more viral-host and plasmid-host associations in metaMDBG (333 viral-host and 32 plasmid-host associations) than hifiasm-meta (123 viral-host and 25 plasmid-host associations). Taken together, these results indicate that we consistently detected more viral sequences in the metaMDBG contigs, but more proviral sequences in the hifiasm-meta contigs.

## DISCUSSION

Cataloging the microbial diversity of the human gut presents major challenges, and large numbers of species and strains remain unknown. HiFi metagenomic sequencing provides a rapid and scalable alternative to culturomics, and also offers substantial improvements over short-read sequencing. Here, we performed a deep-sequencing experiment on a pooled human gut microbiome - a challenging sample containing an inflated number of species and strains. We generated large numbers of MAGs using different combinations of long-read assembly, binning, and consolidation methods (Fig. 2). Our study demonstrates that metagenome assembly with HiFi reads can produce large numbers of highly complete MAGs, corroborating the findings of previous studies (Bickhart et al. 2022; Feng et al. 2022; Kato et al. 2022; Kim et al. 2022; Zhang et al. 2022; Benoit et al. 2023; Jiang et al. 2023; Saak et al. 2023; Schaerer et al. 2023; Masuda et al. 2024). Across all method combinations, we found up to 595 total MAGs, 277 HQ-MAGs, and 98 single-contig HQ-MAGs (including 70 circular). These MAGs represent up to 455 species clusters (Fig. 4), with 15% of the MAGs representing uncultured species (including >35 HQ-MAGs). Based on our results, we discuss the relative performance and trade-offs for different methods, which overall we find to be complementary.

We used bioinformatic and proximity ligation binning methods to group contigs into putative MAGs, and developed a new framework to explicitly compare MAGs across binning strategies (Fig. 1). From a simple performance perspective, we found ProxiMeta generally produced more total MAGs than the HiFi-MAG-Pipeline (Fig. 2, Supplementary Figs. S3, S4). However, our framework with pb-MAG-mirror allowed us to move beyond counting MAGs to better understand how the contents of MAGs were distributed across both methods. In an ideal scenario, two orthogonal methods would produce MAG sets with a large proportion of highly similar MAGs (e.g., MAGs with near-identical contig contents), increasing our confidence in the biological reality of those genomes. In the worst case scenario, the methods could produce MAG sets with no sensible overlap - the contig contents of individual MAGs are highly mixed across MAGs in the other method. It is also possible to recover unique MAGs (e.g., those with contig contents not appearing in the other method), which may highlight strengths or blindspots for a given method. In our study, we found ∼65% of total MAGs were classified as highly similar across the binning methods, with ∼20% being highly mixed and another ∼15% being unique to each method (Fig. 2). These proportions were consistent across hifiasm-meta and metaMDBG, suggesting they are robust to the starting contigs. The large proportion of highly similar MAGs is reassuring, as the same core set of MAGs (roughly ∼320) can be recovered using HiFi-MAG-Pipeline or ProxiMeta (Fig. 2). The 20% of highly mixed MAGs (∼100) is somewhat disconcerting, highlighting major disagreements across the binning methods. Furthermore, to perform consolidation a decision must be made about which MAGs are more accurate. As previous studies have demonstrated higher accuracy for bins using proximity ligation information (Burton et al. 2014; Ho et al. 2023), we selected these in our consolidation step.

Finally, we found each method produced ∼15% unique bins each (ranging from 55–100 MAGs), highlighting information captured by only one method. Unique bins from HiFi-MAG-Pipeline had significantly lower coverages than those from ProxiMeta (which requires a minimum coverage threshold), suggesting bioinformatic binning may perform better at recovering very low coverage MAGs. Conversely, ProxiMeta can use contact information to associate contigs and refine bins, potentially grouping contigs otherwise missed by tetranucleotide frequencies and coverage distributions. By comparing and consolidating the MAGs from ProxiMeta and HiFi-MAG-Pipeline, we identified overlap and incorporated complementary aspects of both methods to improve MAG yields. For example, ProxiMeta provided a 3–8% increase in total MAGs over the HiFi-MAG-Pipeline, but pb-MAG-mirror resulted in an 11–22% increase in total MAGs over both binning methods. Overall, our comparison revealed that high numbers of MAGs can be obtained using both binning methods, a large core set of MAGs is shared across the methods, and that consolidation can incorporate the unique information captured by each.

We used hifiasm-meta and metaMDBG for metagenome assembly, and based on performance we found different strengths for each method. At the highest level, we were interested in which assembly method produced the most MAGs. In general, hifiasm-meta produced more total MAGs than metaMDBG (Fig. 2), with some exceptions in the downsampled datasets. However, regarding genome quality we found that metaMDBG consistently produced more HQ-MAGs than hifiasm-meta (Figs. 2, 3). Specifically, HQ-MAGs make up 50% of the total MAGs for metaMDBG in the consolidated set, but only 30% of the total for hifiasm-meta. We also investigated how well each method assembled strain-level variation. Here, hifiasm-meta produced strains for 56% more species and assembled 57% more total strain-level MAGs relative to metaMDBG, for the consolidated MAGs (Fig. 4, Table 3). Of the species with strains assembled, we wanted to understand whether strain-level variation was represented as two or more near-complete genomes (e.g., HQ-MAGs), as one near-complete genome and several incomplete genomes, or as many fragmented genomes. Despite their differences in the total number of strains assembled, hifiasm-meta and metaMDBG recovered two or more strain-level, near-complete genomes for a similar number of species in the consolidated MAGs (21 and 24, respectively; Supplementary Table S5). For both assembly methods, strains were most often represented by one near-complete genome and multiple partial genomes. We observed that for hifiasm-meta, strains were also commonly represented by sets of partial genomes, whereas this was infrequent for metaMDBG. These results highlight that both assembly methods are capable of assembling strain-level variation into two or more near-complete genomes, resolving strains remains challenging, and that hifiasm-meta currently produces more strain-level information.

We investigated how many MAGs were shared across the assembly methods in the consolidated sets. We used strict criteria to identify unequivocal matches across the assembly methods, which resulted in 125 high-confidence matches at the strain level (Fig. 6). These unequivocally shared MAGs tended to be very high-quality, with greater completeness scores, fewer contigs (including many single-contig), and higher depth of coverage relative to MAGs not shared across the methods (Fig. 6). Although this only represents ∼22% of the total MAGs in each method, we emphasize that even slight differences in the assembly or binning of contigs could disqualify a potential match. It is also possible that partial strain-collapsing in some MAGs could impact this type of analysis, which would be exacerbated in this high-strain-diversity sample. Remarkably, there are 17 matched MAGs which both consist of a single-contig, are within 200 bp of the total length of one another (across a 1.42–6.09 Mb size range), and display only 2–100 nucleotide differences (Supplementary Table S10). These results highlight that the use of highly accurate long reads can lead to the recovery of nearly identical MAGs, despite the use of disparate assembly and binning methods.

Beyond bacterial and archaeal genomes, viral and plasmid genomes pose a different set of assembly challenges (Antipov et al. 2020). For long reads specifically, their size range can fall below the length of a typical HiFi read (10–20 kb), and their structure is quite variable. We used ProxiMeta, VirSorter2, and geNomad to identify and annotate viral sequences, and in most cases recovered more total viral sequences for metaMDBG (Supplementary Table S11). With the ProxiMeta platform, we detected more viral and plasmid host associations for metaMDBG versus hifiasm-meta. These results indicate metaMDBG may have an advantage for assembling smaller, viral sequences, but that additional work is required to improve assembly for these highly variable genomes.

The routine production of single-contig HQ-MAGs from long-read metagenome assemblies is a recent phenomenon, and we propose it requires some important considerations. First, a majority of binning algorithms operate with an implicit assumption that the assembled contigs represent fragmented genomes, and this assumption is often violated with HiFi metagenome assembly. Performing naïve binning on HiFi assemblies can result in the mis-binning of single-contig MAGs with additional contigs, inflating their contamination score and causing removal during quality filtering (Feng et al. 2022; Benoit et al. 2023). For this reason, we built a “completeness-aware” strategy into the HiFi-MAG-Pipeline, and propose it is an essential step for any long-read binning workflow. This strategy involves screening all contigs longer than 500kb to obtain completeness and contamination scores, and moving all highly complete, single-contig MAGs into individual bins. The subsequent binning steps therefore only include incomplete contigs, preventing the mis-binning and removal of any SC-HQ-MAGs.

Second, the quality standards proposed by the Genomic Standards Consortium (Bowers et al. 2017) have proven useful for comparing short-read MAGs, but they do not capture additional information relevant for long-read MAGs. Specifically, contiguity (i.e., the number of contigs per MAG) is becoming increasingly important for evaluating quality. For example, a short-read MAG can be defined as high-quality based on the presence of single-copy genes despite containing hundreds of contigs (and potentially missing large portions of the accessory genome). An ideal HQ-MAG is composed of a single contig per chromosome, and these are often distinguished in long-read studies (Bickhart et al. 2022; Feng et al. 2022; Kim et al. 2022; Zhang et al. 2022; Benoit et al. 2023; Jiang et al. 2023; Masuda et al. 2024), including ours (labeled as SC-HQ-MAGs). However, there is currently no distinction between HQ-MAGs composed of a few long contigs (e.g., hundreds of kilobases, or megabases) versus those containing hundreds of short contigs (e.g., tens of kilobases or shorter), despite the former being much higher quality.

We propose this idea should be reflected in future terminology, allowing distinctions to be made along the contiguity spectrum. Beyond contiguity, circularity is sometimes also highlighted as an important quality metric (Bickhart et al. 2022; Kim et al. 2022; Zhang et al. 2022; Jiang et al. 2023; Masuda et al. 2024). However, Kim et al. (2022) demonstrated that circular MAGs can contain large gaps, and circularity alone should not be taken as a reliable indicator of completeness. Furthermore, linear contigs can sometimes be closed using additional read-mapping information. Therefore, information about linear vs. circular SC-HQ-MAGs can be useful, but may not be as meaningful as other quality categories. Although defining new standards is beyond the scope of this study, we suggest these criteria should be re-evaluated by the community as long-read metagenomics continues to become more mainstream.

## METHODS

### Sample

The ZymoBIOMICS™ Fecal Reference with TruMatrix™ Technology [D6323] (Zymo Research – CA, USA) was made by collecting multiple fecal samples from healthy human volunteers following ethical guidelines and informed consent procedures. Samples were collected using sterile collection tubes and stored at −80°C until processing. Collections from multiple donors were pooled and homogenized in one large suspension together with the microbiome preservative reagent DNA/RNA Shield™ (Zymo Research – CA, USA) to prevent outgrowths or depletions of microbial taxa. The fecal suspension was then distributed to containers to be stored at −80°C until aliquoting to individual tubes. Each stored container was tested individually by sequencing to confirm consistency prior to aliquoting to individual tubes. All aliquots have been demonstrated to have consistent metagenomic and metatranscriptomic profiles.

### Sequencing

DNA was extracted using the ZymoBIOMICS DNA Miniprep Kit (D4300), which involved either mechanical (Vortex Genie 2) or enzymatic lysis. Fragment sizes were measured using a Femto Pulse (Agilent). A total of eight SMRTbell libraries were prepared from four DNA extractions. Libraries for the PacBio Sequel IIe system were created using the SMRTbell Express Template Prep Kit 2.0 (n=3) or SMRTbell Prep Kit 3.0 (n=3), and libraries were prepared for the Revio system using the SMRTbell Prep Kit 3.0 (n=2). Size selection was performed using 3.7X 35% Ampure SPRIselect (Beckman Coulter), which targeted the removal of fragments <3kb. Six libraries were sequenced on the Sequel IIe system (using 40–60 ng library input) and two on the Revio system (using 130–140 ng library input), with each library sequenced on an individual 8M or 25M SMRT Cell, respectively. For the Sequel IIe system, HiFi reads were generated using ccs v6.2, whereas for the Revio system HiFi reads were generated on-instrument using Google DeepConsensus (Baid et al. 2023). For downstream data analysis, we combined the HiFi data obtained from Sequel IIe and Revio. To investigate the effects of lower data levels on assembly, we performed downsampling on the Sequel IIe dataset. Our downsampling design includes the equivalent of six 8M SMRT Cells down to one cell, along with a 2plex, 4plex, and 8plex on one cell (Supplemental Table S1).

A proximity ligation (Hi-C) library was prepared from an aliquot of the Fecal Reference standard using the ProxiMeta Hi-C v4.0 Kit from Phase Genomics according to the manufacturer provided protocol (Lieberman-Aiden et al. 2009). Briefly, intact cells were crosslinked using a formaldehyde solution, simultaneously digested using the *Sau3AI* and *MlucI* restriction enzymes, and proximity ligated with biotinylated nucleotides to create chimeric molecules composed of fragments from different regions of microbial genomes that were physically proximal *in vivo*. Proximity ligated DNA molecules were pulled down with streptavidin beads and processed into an Illumina-compatible sequencing library. Sequencing was performed on an Illumina NovaSeq with PE150 read pairs, producing 181.7M reads in total.

### Metagenome assembly and binning

We performed assemblies with hifiasm-meta r74 (Feng et al. 2022) and metaMDBG v0.3 (Benoit et al. 2023) using default parameter settings. The resulting contigs were processed using two distinct methodologies, including bioinformatic binning and proximity ligation (Hi-C) binning. A visual overview of our analysis workflow is shown in Figure 1.

For bioinformatic binning, we developed a new version of the HiFi-MAG-Pipeline (available from: https://github.com/PacificBiosciences/pb-metagenomics-tools), which uses a completeness-aware binning strategy. The workflow begins by identifying all contigs longer than 500kb which display >90% completeness and <10% contamination, as measured by CheckM2 v1.0.1 (Chklovski et al. 2023). These highly complete, single-contig MAGs are removed from the contig set and placed in individual bins. The remaining set of incomplete contigs is then subjected to binning using the long-read mode of SemiBin2 v1.5 (Pan et al. 2022, 2023) and with MetaBAT2 v2.15 (Kang et al. 2019). To obtain coverage scores per contig, minimap2 v2.17 (Li 2018, 2021) is used to map reads to the contig set, and the jgi_summarize_bam_contig_depths script of MetaBAT2 is used to summarize depth per contig. Settings for MetaBAT2 included -m 30000, whereas SemiBin2 used the single_easy_bin module and included the --self-supervised, --sequencing-type=long_reads, and --environment=human_gut flags. The two bin sets are de-replicated using DAS_Tool v1.1.6 (Sieber et al. 2018), and the de-replicated bin set is assessed with CheckM2. The bins which pass minimum criteria for single-copy gene (SCG) completeness and contamination are added to the set of highly complete, single-contig MAGs identified in the first step. Finally, the Genome Taxonomy Database Toolkit (GTDB-Tk) v2.1.1 (Chaumeil et al. 2019, 2022) is used to assign taxonomy to all filtered MAGs.

MAGs were categorized as medium-quality or high-quality (MQ-MAG, HQ-MAG, respectively): HQ-MAGs require ≥90% SCG completeness and ≤5% contamination, whereas MQ-MAGs fall below these criteria but display ≥50% SCG completeness and ≤10% contamination. We note that the total number of MAGs reported in our results is equal to the number of HQ-MAGs plus MQ-MAGs. We also include a third category for the highest level of quality, which we label as a single-contig, high-quality MAG (SC-HQ-MAG). This category includes the same criteria as an HQ-MAG, plus the presence of a single contig (circularity optional).

In addition to bioinformatic binning, we used proximity ligation to generate bins (Press et al. 2017). Proximity ligation sequencing files and assembled contigs were uploaded to the cloud-based ProxiMeta platform (Uritskiy et al. 2021). Proximity ligation reads were aligned using BWA-MEM (Li & Durbin 2010) with the −5SP options specified, and all other options default. SAMBLASTER (Faust & Hall 2014) was used to flag PCR duplicates, which were later excluded from analysis. Alignments were then filtered with samtools (Li et al. 2009) using the -F 2304 filtering flag to remove non-primary and secondary alignments. Metagenome deconvolution was performed with ProxiMeta (Press et al. 2017; Stewart et al. 2018), resulting in the creation of putative genome and genome fragment clusters, as well as viral, plasmid, and AMR gene host annotations. For consistency, we ran the bins inferred by the ProxiMeta workflow through CheckM2, filtered based on the above quality criteria, and assigned taxonomy using GTDB-Tk v2.1.1. This allowed a more direct comparison to the bins recovered by HiFi-MAG-Pipeline.

### Binning method comparison

We compared the HiFi-MAG-Pipeline and ProxiMeta binning results using a new approach we developed, called pb-MAG-mirror. We compared the contig content of MAGs across methods to identify four categories: identical bins, superset/subset bins, mixed bins, and unique bins (Fig. 1). Identical bins occur when a bin from each method contains the exact same contig set. A subset bin occurs when the contig set of a bin is fully contained in a bin of the alternate method (i.e., the superset bin). We further require that any additional contigs in the superset bin do not occur in any other bin (e.g., any additional contigs present in the superset bin were unique to the superset bin). Lastly, a unique bin occurs when it contains a set of contigs that does not occur in any bin of the alternative method. If a single contig of the set can be found in a bin of the alternative method, the bin cannot be classified as unique. All bins not falling into these categories (identical, subset/superset, unique) are considered mixed, as they contain two or more contigs that occur in two or more bins of the alternative method. For these mixed bins, we perform cross-method pairwise comparisons. We examine the shared contig content of each comparison to identify the best match, which is selected based on the percentage of total shared bases. We consider high-similarity (HS) matches as those with ≥80% shared bases in each bin, medium-similarity (MS) as those with ≥50% shared bases in each bin, and low-similarity (LS) as those with <50% shared bases in each bin. Based on the category results, pb-MAG-mirror can be used to consolidate the two bin sets. The default behavior is to include one representative bin for each of the identical bins, all superset bins and unique bins from both sets, and for cross-set mixed matches the bin with the highest completeness score is selected. However, for cross-set mixed matches the bins from one set can be preferentially selected instead. For our comparisons, for the cross-set mixed matches we selected ProxiMeta bins. We refer to the resulting MAGs as the consolidated MAG set (Fig. 1). To identify potential differences in characteristics of unique bins between HiFi-MAG-Pipeline and ProxiMeta, we compared completeness scores, number of contigs, and average depth of coverage for the unique bins. pb-MAG-mirror is available as an open-source workflow at: https://github.com/PacificBiosciences/pb-metagenomics-tools.

### Taxonomic and phylogenetic analyses

We quantified the number of species contained in the various MAG sets using two approaches. First, we used the taxonomy assigned by GTDB-Tk to count the number of unique species assignments. However, several genomes were not assigned to the species level. These unassigned genomes could represent one or more species groups. To address this issue, we used dRep v3.4.3 (Olm et al. 2017) to cluster MAGs into species-level groups based on an average nucleotide identity (ANI) of 95% (Jain et al. 2018; Olm et al. 2020). The default settings were used, which involved the ANImf algorithm for genome comparisons, 90% ANI for creating primary clusters, and 95% ANI for creating secondary clusters. We regarded each resulting cluster as a unique species. For strain-level counts, we aggregated counts based on genomes with the same species (GTDB-TK) or cluster (dRep) assignment.

We reconstructed the phylogenetic relationships of the MAGs from the consolidated MAG sets for each assembly method. We obtained protein alignments for core bacterial and archaeal genes (Pfam, TIGRFAM) using the identify and align modules of GTDB-Tk. The core gene sets differ in number between bacteria and archaea (n=120 and n=53, respectively), with five genes shared across both sets. We created a concatenated supermatrix of all genes and analyzed the supermatrix using RAxML v8.2.12 (Stamatakis 2014). Tree searches used the gamma model of rate heterogeneity and JTTDCMUT amino acid substitution model. We visualized the phylogeny using the Interactive Tree of Life (iTOL) v6.8.1 online tool (Letunic & Bork 2021).

### Comparison of assembly methods

We compared contigs from hifiasm-meta and metaMDBG using two approaches. First, we aligned the contig sets using minimap2, and calculated the percent of bases covered based on 98 or 99% identity. Second, for contigs >500kb we calculated Mash similarities (MinHash) using mash (Ondov et al. 2016) and FracMinHash-based similarities using sourmash (Pierce et al. 2019; Irber et al. 2022). There were 996 contigs >500kb for hifiasm-meta and 867 for metaMDBG. Concordantly high values for both Mash and FracMinHash similarities were taken as strong evidence for a contig match across assemblers, whereas a high value for Mash and lower value for FracMinHash indicated a contig was likely contained within the other contig. A range of minimum cutoffs for each metric was explored.

We also investigated the number of HiFi reads that could be reliably mapped to the contigs or consolidated MAGs, per assembly method. We performed read mapping using minimap2 with HiFi settings (-x map-hifi -k19 -w19 --secondary=no), and subsequently filtered the alignments. We kept primary alignments in which ≥90% of the HiFi read was aligned to the reference, and with ≥95% identity (defined as number of matched query bases divided by total query bases in the alignment). To identify potential differences in MAG characteristics across the assembly methods, we compared the average depth of coverage and number of contigs for the consolidated HQ- and MQ-MAGs.

In an effort to understand how representative the MAGs are of the total species diversity in the sample, we compared 16S rRNA sequences obtained from the HiFi reads with those contained in MAGs (Feng & Li 2024). Methods were highly similar to those described in Feng and Li (2024). In brief, HiFi reads containing 16S regions were discovered by mapping to the SILVA database (Quast et al. 2013), rRNA genes were identified using barrnap v0.9 (https://github.com/tseemann/barrnap), annotated using the RDP classifier (Cole et al. 2014), and OTUs were defined using greedy incremental clustering and 99% mismatch identity. The MAG 16S sequences were identified and extracted from the consolidated MAG sets from hifiasm-meta and metaMDBG, and mapped to the OTU set from the HiFi reads. A MAG could have more than one OTU assignment if it contained multiple, distinct 16S copies.

We sought to determine the number of identical or nearly-identical MAGs that were unequivocally shared across the assembly methods. To accomplish this, we estimated the intersection of the consolidated MAG sets from each assembly method, using minimum thresholds for aligned sequence lengths and ANI. We used dRep to generate clusters for all consolidated MAGs from hifiasm-meta and metaDMBG, using the ANImf algorithm with 90% ANI for primary clusters and 99% ANI for secondary clusters. For each resulting cluster, we first determined if it contained at least one MAG per assembly method. If so, we then performed pairwise comparisons of all MAGs from hifiasm-meta to metaMDBG to identify unequivocal matches. To understand the effects of parameter values on the number of matches, we explored using 95% or 99% ANI and a range of values for the minimum percent of bases per MAG required (70–100%). Based on these results, we defined an unequivocal match as requiring ≥90% of the total bases to be aligned (per genome) with ≥99% ANI (e.g., a nearly identical match at the strain level)

### Mobile element association and viral annotation

Mobile element host-association was performed using the cloud-based ProxiMeta platform (Uritskiy et al. 2021). Briefly, viral contigs were identified using VIBRANT (Kieft et al. 2020), and plasmid contigs were identified using BLAST and the NCBI Plasmid Database. Alignments of long-range Hi-C linkage data were used to identify viral-host and plasmid-host linkages. A combination of the Hi-C link count, mobile element read depth, and MAG read depth were then used to estimate the average copy count of each mobile element in each MAG. The density of Hi-C links per kb^2^ of sequence between the mobile element and the MAG was then compared to the connectivity of the MAG to itself and normalized to the estimated copy count to compute the normalized connectivity ratio. Mobile-host linkages were then filtered to keep only connections with at least 2 Hi-C read links between the mobile and host MAG, a connectivity ratio of 0.1, and intra-MAG connectivity of 10 links to remove false positives. For the final threshold value, a receiver operating characteristic (ROC) curve is used to determine the optimal copy count cut-off value, which is the value that removes the maximum number of virus-host links while still finding at least one host for the maximum number of mobile elements. Additional filtering removed linkages with an average copy count less than 80% of the highest copy count value for the given mobile element sequence.

As an alternative to ProxiMeta, we also predicted viral and proviral sequences using geNomad (Camarago et al. 2023) and VirSorter2 (Guo et al. 2021). geNomad (ver 1.8.0) was run in “relaxed” mode. VirSorter2 (v2.2.4) was run with all steps and results were verified with CheckV (v1.0.1) using “end-to-end” mode (Nayfach et al. 2021). VirSorter2 viral counts include all quality types reported from CheckV (complete, high quality, medium quality, low quality, and not determined).

## DATA ACCESS

HiFi sequencing data are publicly available on NCBI (PRJNA1110296) and from: https://downloads.pacbcloud.com/public/revio/2023Q3/ZymoTrumatrix/ https://downloads.pacbcloud.com/public/sequelii/2023Q3/ZymoTrumatrix/

The contigs, MAGs, and add sets for each binning (HiFi-MAG-Pipeline, ProxiMeta, consolidated) and assembly (hifiasm-meta, metaMDBG) combination are publicly available on the Open Science Framework: https://osf.io/cwqzr/

pb-MAG-mirror is publicly available at: https://github.com/PacificBiosciences/pb-metagenomics-tools

## ACKNOWLEDGMENTS

CQ acknowledges the support of the Biotechnology and Biological Sciences Research Council (BBSRC), part of UK Research and Innovation; Earlham Institute Strategic Programme Grant (Decoding Biodiversity) BBX011089/1 and its constituent work package BBS/E/ER/230002C; the Core Strategic Programme Grant (Genomes to Food Security) BB/CSP1720/1 and its constituent work packages BBS/E/T/000PR9818 and BBS/E/T/000PR9817; and the Core Capability Grant BB/CCG2220/1. GB was supported by the ERC IndexThePlanet grant (UE ERC-2022-COG INDEXTHEPLANET). Work by members of Phase Genomics were supported by NIH/NIAID grants R44AI172703 and R44AI162570 as well as funding from the Bill and Melinda Gates Foundation.

