## Supplemental File for "Highly accurate metagenome-assembled genomes from human gut microbiota using long-read assembly, binning, and consolidation methods"


| **Description** | **Page** |
|---|---|
| Table S1. pb-MAG-mirror category counts | 2 |
| Table S2. Summary of bioinformatically downsampled HiFi sequencing datasets | 2 |
| Table S3. MAG counts for downsampled datasets | 3 |
| Table S4. Logarithmic trendlines describing total data versus consolidated MAG yield | 5 |
| Table S5. Counts of dRep species clusters containing different numbers of strains | 5 |
| Table S6. Counts of strains for species displaying 4 or more strain-level MAGs | 6 |
| Table S7. Taxonomic lineages and counts of HQ-MAGs unassigned at the species rank | 7 |
| Table S8. Mash and FracMinHash comparisons for contigs >500kb | 8 |
| Table S9. Number and percent of reads aligned to contigs or consolidated MAG sets | 9 |
| Table S10. Summary of 125 unequivocal MAG matches | 10 |
| Table S11. Mobile element counts | 15 |
| Figure S1. HiFi read characteristics between Sequel IIe and Revio | 16 |
| Figure S2. MAG characteristics by pb-MAG-mirror categories | 17 |
| Figure S3. MAG counts for hifiasm-meta across all downsampled datasets | 18 |
| Figure S4. MAG counts for metaMDBG across all downsampled datasets | 19 |
| Figure S5. Parameter comparisons for unequivocally matched MAGs | 20 |
| Figure S6. Overlap of viral contigs across assemblers | 21 |

**Supplementary Table S1.** Summary of MAG comparison analyses for the three main datasets.

| Assembly method | Binning method | Total MAGs | Identical | Superset | Subset | Mixed-HS | Mixed-MS | Mixed-LS | Unique |
|---|---|---|---|---|---|---|---|---|---|
| hifiasm-meta | HiFi-MAG-Pipeline | 485 | 148 | 27 | 103 | 44 | 55 | 35 | 73 |
| | ProxiMeta | 522 | 148 | 103 | 27 | 44 | 55 | 39 | 106 |
| metaMDBG | HiFi-MAG-Pipeline | 480 | 116 | 26 | 122 | 53 | 76 | 32 | 55 |
| | ProxiMeta | 492 | 116 | 122 | 26 | 53 | 76 | 34 | 65 |

Mixed MAGs include high-similarity (HS; >=80% shared bases), medium-similarity (>=50% shared bases), and low-similarity (LS, <50% shared bases).

**Supplementary Table S2.** Summary of the bioinformatically downsampled HiFi sequencing datasets.

| Dataset | Downsampled sequencing | HiFi reads (million) | HiFi yield (Gb) | Mean read length | Mean QV |
|---|---|---|---|---|---|
| SQIIe 6Cell | 6 SMRT Cells (8M) | 17.15 | 120.85 | 7,046 | 40.2 |
| SQIIe 5Cell | 5 SMRT Cells (8M) | 14.32 | 102.87 | 7,180 | 40.2 |
| SQIIe 4Cell | 4 SMRT Cells (8M) | 11.89 | 88.68 | 7,453 | 40.2 |
| SQIIe 3Cell | 3 SMRT Cells (8M) | 8.56 | 68.12 | 7,958 | 40.2 |
| SQIIe 2Cell | 2 SMRT Cells (8M) | 6.25 | 53.49 | 8,547 | 40.2 |
| SQIIe 1Cell | 1 SMRT Cell (8M) | 2.92 | 25 | 8,548 | 40.2 |
| SQIIe 2plex | 2plex on 1 SMRT Cells (8M) | 1.46 | 12.51 | 8,551 | 40.2 |
| SQIIe 4plex | 4plex on 1 SMRT Cells (8M) | 0.73 | 6.25 | 8,550 | 40.2 |
| SQIIe 8plex | 8plex on 1 SMRT Cells (8M) | 0.36 | 3.12 | 8,548 | 40.2 |

**Supplementary Table S3.** Counts of single-contig high-quality (SC-HQ) MAGs, high-quality (HQ) MAGs, and medium-quality (MQ) MAGs for each dataset and binning method combination.

| Sample | Assembly method | Binning method | HQ-MAGs | MQ-MAGs | Total |
|---|---|---|---|---|---|
| SQIIe 6Cell | hifiasm-meta | HiFi-MAG-Pipeline | 106 | 247 | 353 |
| | | ProxiMeta | 116 | 277 | 393 |
| | | Consolidated | 128 | 321 | 449 |
| | metaMDBG | HiFi-MAG-Pipeline | 174 | 188 | 362 |
| | | ProxiMeta | 186 | 204 | 390 |
| | | Consolidated | 200 | 241 | 441 |
| SQIIe 5Cell | hifiasm-meta | HiFi-MAG-Pipeline | 101 | 221 | 322 |
| | | ProxiMeta | 113 | 291 | 404 |
| | | Consolidated | 119 | 322 | 441 |
| | metaMDBG | HiFi-MAG-Pipeline | 171 | 178 | 349 |
| | | ProxiMeta | 194 | 168 | 362 |
| | | Consolidated | 211 | 194 | 405 |
| SQIIe 4Cell | hifiasm-meta | HiFi-MAG-Pipeline | 111 | 195 | 306 |
| | | ProxiMeta | 118 | 230 | 348 |
| | | Consolidated | 125 | 265 | 390 |
| | metaMDBG | HiFi-MAG-Pipeline | 167 | 177 | 344 |
| | | ProxiMeta | 185 | 171 | 356 |
| | | Consolidated | 199 | 200 | 399 |
| SQIIe 3Cell | hifiasm-meta | HiFi-MAG-Pipeline | 118 | 182 | 300 |
| | | ProxiMeta | 132 | 225 | 357 |
| | | Consolidated | 136 | 259 | 395 |
| | metaMDBG | HiFi-MAG-Pipeline | 153 | 162 | 315 |
| | | ProxiMeta | 159 | 145 | 304 |
| | | Consolidated | 174 | 178 | 352 |
| SQIIe 2Cell | hifiasm-meta | HiFi-MAG-Pipeline | 106 | 176 | 282 |
| | | ProxiMeta | 122 | 230 | 352 |
| | | Consolidated | 127 | 250 | 377 |
| | metaMDBG | HiFi-MAG-Pipeline | 138 | 145 | 283 |
| | | ProxiMeta | 155 | 149 | 304 |

| | | | | | |
|---|---|---|---|---|---|
| | | Consolidated | 162 | 173 | 335 |
| SQIIe 1Cell | hifiasm-meta | HiFi-MAG-Pipeline | 71 | 143 | 214 |
| | | ProxiMeta | 83 | 191 | 274 |
| | | Consolidated | 86 | 203 | 289 |
| | metaMDBG | HiFi-MAG-Pipeline | 95 | 115 | 210 |
| | | ProxiMeta | 93 | 109 | 202 |
| | | Consolidated | 108 | 135 | 243 |
| SQIIe 2plex | hifiasm-meta | HiFi-MAG-Pipeline | 39 | 88 | 127 |
| | | ProxiMeta | 40 | 131 | 171 |
| | | Consolidated | 44 | 139 | 183 |
| | metaMDBG | HiFi-MAG-Pipeline | 66 | 67 | 133 |
| | | ProxiMeta | 62 | 74 | 136 |
| | | Consolidated | 70 | 89 | 159 |
| SQIIe 4plex | hifiasm-meta | HiFi-MAG-Pipeline | 21 | 73 | 94 |
| | | ProxiMeta | 18 | 84 | 102 |
| | | Consolidated | 25 | 96 | 121 |
| | metaMDBG | HiFi-MAG-Pipeline | 31 | 57 | 88 |
| | | ProxiMeta | 31 | 50 | 81 |
| | | Consolidated | 37 | 71 | 108 |
| SQIIe 8plex | hifiasm-meta | HiFi-MAG-Pipeline | 7 | 52 | 59 |
| | | ProxiMeta | 8 | 50 | 58 |
| | | Consolidated | 10 | 63 | 73 |
| | metaMDBG | HiFi-MAG-Pipeline | 7 | 32 | 39 |
| | | ProxiMeta | 6 | 33 | 39 |
| | | Consolidated | 9 | 45 | 54 |

HQ-MAGs require ≥90% single-copy genes (SCG) completeness and ≤5% contamination, whereas MQ-MAGs fall below the HQ thresholds but display ≥50% SCG completeness and ≤10% contamination.

**Supplementary Table S4.** Summary of logarithmic trendlines describing total data versus consolidated MAG yield.

| **Assembly method** | **MAG category** | **Equation** | **$R^2$** |
|---|---|---|---|
| hifiasm-meta | HQ-MAGs | y = 38.581ln(x) - 41.058 | 0.9472 |
| | MQ-MAGs | y = 76.302ln(x) - 44.251 | 0.9666 |
| metaMDBG | HQ-MAGs | y = 58.758ln(x) - 70.777 | 0.9799 |
| | MQ-MAGs | y = 51.354ln(x) - 26.899 | 0.9719 |

**Supplementary Table S5.** Counts of dRep species clusters containing different numbers of strains, for the consolidated MAG sets.

| Assembly method | Criterion | 1 strain | 2 strains | 3 strains | 4 strains | 5 strains |
|---|---|---|---|---|---|---|
| hifiasm-meta | All | 349 | 68 | 20 | 10 | 2 |
| | ≥1 HQ-MAG | 108 | 33 | 13 | 6 | 1 |
| | ≥2 HQ-MAGs | - | 13 | 7 | 1 | 0 |
| | ≥3 HQ-MAGs | - | - | 3 | 0 | 0 |
| metaMDBG | All | 391 | 46 | 10 | 6 | 2 |
| | ≥1 HQ-MAG | 244 | 34 | 8 | 4 | 1 |
| | ≥2 HQ-MAGs | - | 18 | 4 | 1 | 1 |
| | ≥3 HQ-MAGs | - | - | 2 | 0 | 1 |

**Supplementary Table S6**. Counts of strains for the most common species displaying 4 or more strain-level MAGs in at least one assembly method.

| **Species** | **hifiasm-meta strains** | **metaMDBG strains** |
|---|---|---|
| *Adlercreutzia celatus_A / equolifaciens* | **4** | 5 |
| *Agathobacter faecis* | **4** | **2** |
| *Agathobaculum butyriciproducens* | **4** | **5** |
| *Bacteroides uniformis* | 4 | **4** |
| *Blautia_A massiliensis* | **4** | **1** |
| *CAG-41 sp900066215* | **3** | 4 |
| *Copromonas sp000435795* | **4** | **2** |
| *Dorea_A longicatena* | 4 | 1 |
| *Dysosmobacter sp001916835* | 2 | **4** |
| *Faecalibacterium prausnitzii_D* | 5 | 3 |
| *Faecalibacterium prausnitzii_G* | **5** | **4** |
| *Faecalibacterium sp900539945* | 2 | **4** |
| *Lachnospira eligens_A / sp003451515* | 4 | 4 |
| *Phocaeicola dorei/vulgatus* | 4 | **3** |
| *Ruminococcus_B gnavus* | **4** | **2** |

Numbers in bold indicate ≥1 HQ-MAG present, and underline indicates ≥2 HQ-MAGs in the strain set.

**Supplemental Table S7.** Taxonomic lineages and counts of HQ-MAGs that were unassigned at the species rank.

| **Taxa** | **hifiasm-meta Consolidated** | **metaMDBG Consolidated** |
|---|---|---|
| d__Bacteria;p__Actinobacteriota;c__Coriobacteriia;o__Coriobacteriales;f__Coriobacteriaceae;g__Collinsella;s__ | 1 | 1 |
| d__Bacteria;p__Actinobacteriota;c__Coriobacteriia;o__Coriobacteriales;f__Eggerthellaceae;g__CAAEEV01;s__ | 1 | 1 |
| d__Bacteria;p__Desulfobacterota_I;c__Desulfovibrionia;o__Desulfovibrionales;f__Desulfovibrionaceae;g__Desulfovibrio;s__ | 1 | 1 |
| d__Bacteria;p__Firmicutes_A;c__Clostridia;o__Lachnospirales;f__Lachnospiraceae;g__;s__ | 1 | 1 |
| d__Bacteria;p__Firmicutes_A;c__Clostridia;o__Lachnospirales;f__Lachnospiraceae;g__Blautia_A;s__ | 1 | 1 |
| d__Bacteria;p__Firmicutes_A;c__Clostridia;o__Lachnospirales;f__Lachnospiraceae;g__Eubacterium_G;s__ | 0 | 1 |
| d__Bacteria;p__Firmicutes_A;c__Clostridia;o__Lachnospirales;f__Lachnospiraceae;g__Eubacterium_I;s__ | 0 | 1 |
| d__Bacteria;p__Firmicutes_A;c__Clostridia;o__Lachnospirales;f__Lachnospiraceae;g__Faecalimonas;s__ | 0 | 1 |
| d__Bacteria;p__Firmicutes_A;c__Clostridia;o__Lachnospirales;f__Lachnospiraceae;g__Frisingicoccus;s__ | 1 | 2 |
| d__Bacteria;p__Firmicutes_A;c__Clostridia;o__Lachnospirales;f__Lachnospiraceae;g__JAGZHZ01;s__ | 0 | 1 |
| d__Bacteria;p__Firmicutes_A;c__Clostridia;o__Lachnospirales;f__UBA1390;g__Faecimorpha;s__ | 1 | 1 |
| d__Bacteria;p__Firmicutes_A;c__Clostridia;o__Monoglobales;f__Monoglobaceae;g__UMGS1326;s__ | 1 | 1 |
| d__Bacteria;p__Firmicutes_A;c__Clostridia;o__Oscillospirales;f__Acutalibacteraceae;g__;s__ | 0 | 1 |
| d__Bacteria;p__Firmicutes_A;c__Clostridia;o__Oscillospirales;f__Acutalibacteraceae;g__Avimonas_A;s__ | 0 | 1 |
| d__Bacteria;p__Firmicutes_A;c__Clostridia;o__Oscillospirales;f__Butyricicoccaceae;g__Butyricicoccus_A;s__ | 0 | 1 |
| d__Bacteria;p__Firmicutes_A;c__Clostridia;o__Oscillospirales;f__CAG-272;g__Avispirillum;s__ | 1 | 0 |
| d__Bacteria;p__Firmicutes_A;c__Clostridia;o__Oscillospirales;f__Oscillospiraceae;g__;s__ | 0 | 3 |
| d__Bacteria;p__Firmicutes_A;c__Clostridia;o__Oscillospirales;f__Oscillospiraceae;g__Alloscillospira;s__ | 0 | 1 |
| d__Bacteria;p__Firmicutes_A;c__Clostridia;o__Oscillospirales;f__Oscillospiraceae;g__HGM13006;s__ | 1 | 1 |
| d__Bacteria;p__Firmicutes_A;c__Clostridia;o__Oscillospirales;f__Oscillospiraceae;g__Pseudoscilispira;s__ | 0 | 1 |
| d__Bacteria;p__Firmicutes_A;c__Clostridia;o__Oscillospirales;f__Oscillospiraceae;g__Scatomorpha;s__ | 0 | 1 |
| d__Bacteria;p__Firmicutes_A;c__Clostridia;o__Oscillospirales;f__Oscillospiraceae;g__UMGS1766;s__ | 0 | 2 |
| d__Bacteria;p__Firmicutes_A;c__Clostridia;o__Oscillospirales;f__Ruminococcaceae;g__Gemmiger;s__ | 0 | 1 |
| d__Bacteria;p__Firmicutes_A;c__Clostridia;o__Peptostreptococcales;f__Anaerovoracaceae;g__Alangreenwoodia;s__ | 1 | 1 |
| d__Bacteria;p__Firmicutes_A;c__Clostridia;o__Peptostreptococcales;f__Anaerovoracaceae;g__Copromorpha;s__ | 1 | 1 |
| d__Bacteria;p__Firmicutes_A;c__Clostridia;o__Peptostreptococcales;f__Anaerovoracaceae;g__Emergencia;s__ | 0 | 1 |
| d__Bacteria;p__Firmicutes_A;c__Clostridia;o__RUG12999;f__RUG12999;g__;s__ | 1 | 1 |
| d__Bacteria;p__Firmicutes_A;c__Clostridia;o__UBA1381;f__UBA1381;g__;s__ | 0 | 1 |
| d__Bacteria;p__Firmicutes_B;c__Peptococcia;o__Peptococcales;f__Peptococcaceae;g__UBA7185;s__ | 0 | 1 |

| | | |
|---|---|---|
| d__Bacteria;p__Firmicutes;c__Bacilli;o__Erysipelotrichales;f__Erysipelotrichaceae;g__Traorella;s__ | 0 | 1 |
| d__Bacteria;p__Firmicutes;c__Bacilli;o__RF39;f__UBA660;g__RUG11198;s__ | 1 | 1 |
| d__Bacteria;p__Proteobacteria;c__Alphaproteobacteria;o__RF32;f__CAG-977;g__UBA11549;s__ | 1 | 1 |
| d__Bacteria;p__Proteobacteria;c__Gammaproteobacteria;o__Burkholderiales;f__Burkholderiaceae;g__Oxalobacter;s__ | 1 | 1 |

**Supplemental Table S8.** Results of Mash and FracMinHash comparisons among contigs >500kb.

| Assembly method | Mash | FracMinHash | Match | Contained | Other |
|---|---|---|---|---|---|
| hifiasm-meta | 0.99 | 0.99 | 137 (13.8%) | 131 (13.2%) | 728 (73.1%) |
| | 0.97 | 0.90 | 295 (29.6%) | 315 (31.6%) | 386 (38.8%) |
| | 0.95 | 0.80 | 392 (39.4%) | 444 (44.6%) | 160 (16.1%) |
| | 0.90 | 0.70 | 510 (51.2%) | 474 (47.6%) | 12 (1.2%) |
| metaMDBG | 0.99 | 0.99 | 141 (16.3%) | 125 (14.4%) | 601 (69.3%) |
| | 0.97 | 0.90 | 335 (38.6%) | 245 (28.3%) | 287 (33.1%) |
| | 0.95 | 0.80 | 466 (53.7%) | 290 (33.4%) | 111 (12.8%) |
| | 0.90 | 0.70 | 560 (64.6%) | 291 (33.6%) | 16 (1.8%) |

**Supplemental Table S9.** Number and percent of reads aligned to contigs or consolidated MAG sets.

| Reference | Assembly method | Aligned reads | Percent aligned reads | Filtered aligned reads | Percent filtered aligned reads |
|---|---|---|---|---|---|
| All contigs | hifiasm-meta | 34,495,740 | 99.4 | 27,174,888 | 78.3 |
| | metaMDBG | 34,667,346 | 99.9 | 27,027,899 | 77.9 |
| Consolidated MAGs | hifiasm-meta | 33,590,422 | 96.8 | 21,223,649 | 61.1 |
| | metaMDBG | 33,624,539 | 96.9 | 21,769,453 | 62.7 |
| Viral contigs (excluding provirus) | hifiasm-meta | 17,208,349 | 49.6 | 10,004,402 | 28.8 |
| | metaMDBG | 17,374,140 | 50.0 | 10,505,284 | 30.3 |

A filtered alignment required a primary alignment with ≥90% of the HiFi read aligned with ≥95% nucleotide identity. Total input HiFi reads = 34,715,738.

**Supplemental Table S10.** Summary of 125 unequivocal MAG matches across the consolidated MAG sets from each assembly method.

| Bin1 (hifiasm-meta) | Bin2 (metaMDBG) | Alignment length | Bin1 length | Bin1 alignment coverage | Bin2 length | Bin2 alignment coverage | Similarity errors | ANI |
|---|---|---|---|---|---|---|---|---|
| HiFi-MAG-Pipeline_metabat2.433 | HiFi-MAG-Pipeline_complete.915 | 4,491,940 | 4,479,371 | 1.00 | 4,479,402 | 1.00 | 53 | 0.99999 |
| HiFi-MAG-Pipeline_complete.35 | HiFi-MAG-Pipeline_complete.190 | 1,982,976 | 1,962,913 | 1.01 | 1,962,897 | 1.01 | 49 | 0.99998 |
| ProxiMeta_bin_455 | HiFi-MAG-Pipeline_complete.3 | 1,910,299 | 1,909,439 | 1.00 | 1,993,166 | 0.96 | 85 | 0.99996 |
| HiFi-MAG-Pipeline_complete.279 | HiFi-MAG-Pipeline_complete.327 | 1,784,854 | 1,776,908 | 1.00 | 1,764,850 | 1.01 | 83 | 0.99995 |
| HiFi-MAG-Pipeline_complete.140 | HiFi-MAG-Pipeline_complete.803 | 1,742,160 | 1,742,163 | 1.00 | 1,742,177 | 1.00 | 23 | 0.99999 |
| HiFi-MAG-Pipeline_complete.174 | HiFi-MAG-Pipeline_complete.42 | 1,609,619 | 1,611,148 | 1.00 | 1,609,589 | 1.00 | 11 | 0.99999 |
| HiFi-MAG-Pipeline_complete.315 | HiFi-MAG-Pipeline_metabat2.929 | 1,724,403 | 1,689,538 | 1.02 | 1,708,801 | 1.01 | 121 | 0.99993 |
| HiFi-MAG-Pipeline_semibin2_3 | HiFi-MAG-Pipeline_complete.423 | 2,437,558 | 2,440,106 | 1.00 | 2,432,373 | 1.00 | 25 | 0.99999 |
| HiFi-MAG-Pipeline_complete.227 | HiFi-MAG-Pipeline_complete.216 | 2,013,482 | 2,012,875 | 1.00 | 2,013,293 | 1.00 | 61 | 0.99997 |
| HiFi-MAG-Pipeline_complete.326 | HiFi-MAG-Pipeline_complete.162 | 2,195,297 | 2,195,292 | 1.00 | 2,195,238 | 1.00 | 292 | 0.99987 |
| HiFi-MAG-Pipeline_complete.466 | HiFi-MAG-Pipeline_complete.724 | 2,418,598 | 2,398,609 | 1.01 | 2,398,626 | 1.01 | 71 | 0.99997 |
| **HiFi-MAG-Pipeline_complete.299** | **HiFi-MAG-Pipeline_complete.649** | **3,230,045** | **3,230,054** | **1.00** | **3,230,050** | **1.00** | **10** | **1.00000** |
| HiFi-MAG-Pipeline_complete.569 | HiFi-MAG-Pipeline_complete.912 | 2,930,721 | 2,910,516 | 1.01 | 2,910,530 | 1.01 | 43 | 0.99999 |
| ProxiMeta_bin_56 | ProxiMeta_bin_86 | 3,680,143 | 3,719,984 | 0.99 | 3,677,185 | 1.00 | 1087 | 0.99970 |
| HiFi-MAG-Pipeline_metabat2.343 | HiFi-MAG-Pipeline_semibin2_118 | 3,157,219 | 3,146,048 | 1.00 | 3,233,325 | 0.98 | 214 | 0.99993 |
| ProxiMeta_bin_116 | HiFi-MAG-Pipeline_complete.596 | 3,193,995 | 3,153,999 | 1.01 | 3,149,093 | 1.01 | 355 | 0.99989 |
| **HiFi-MAG-Pipeline_complete.550** | **HiFi-MAG-Pipeline_complete.510** | **2,263,142** | **2,263,158** | **1.00** | **2,263,146** | **1.00** | **8** | **1.00000** |
| HiFi-MAG-Pipeline_complete.575 | ProxiMeta_bin_388 | 2,324,091 | 2,322,461 | 1.00 | 2,341,157 | 0.99 | 89 | 0.99996 |
| HiFi-MAG-Pipeline_complete.342 | HiFi-MAG-Pipeline_complete.11 | 2,492,848 | 2,506,010 | 0.99 | 2,493,457 | 1.00 | 656 | 0.99974 |
| HiFi-MAG-Pipeline_complete.120 | ProxiMeta_bin_473 | 1,899,448 | 1,879,441 | 1.01 | 1,970,983 | 0.96 | 100 | 0.99995 |
| HiFi-MAG-Pipeline_complete.398 | HiFi-MAG-Pipeline_metabat2.47 | 1,963,529 | 1,940,417 | 1.01 | 1,949,019 | 1.01 | 8 | 1.00000 |
| HiFi-MAG-Pipeline_complete.512 | HiFi-MAG-Pipeline_complete.221 | 2,068,430 | 2,067,382 | 1.00 | 2,066,865 | 1.00 | 226 | 0.99989 |
| HiFi-MAG-Pipeline_complete.382 | HiFi-MAG-Pipeline_complete.268 | 2,299,754 | 2,299,765 | 1.00 | 2,301,275 | 1.00 | 3 | 1.00000 |

| | | | | | | | | |
|---|---|---|---|---|---|---|---|---|
| ProxiMeta_bin_301 | ProxiMeta_bin_374 | 2,401,941 | 2,409,965 | 1.00 | 2,415,558 | 0.99 | 7 | 1.00000 |
| HiFi-MAG-Pipeline_complete.20 | HiFi-MAG-Pipeline_complete.351 | 2,391,046 | 2,388,825 | 1.00 | 2,389,806 | 1.00 | 810 | 0.99966 |
| HiFi-MAG-Pipeline_complete.43 | HiFi-MAG-Pipeline_complete.158 | 1,855,430 | 1,856,925 | 1.00 | 1,854,250 | 1.00 | 194 | 0.99990 |
| ProxiMeta_bin_506 | ProxiMeta_bin_524 | 1,563,354 | 1,727,135 | 0.91 | 1,679,723 | 0.93 | 924 | 0.99941 |
| HiFi-MAG-Pipeline_complete.254 | ProxiMeta_bin_421 | 2,207,183 | 2,187,199 | 1.01 | 2,200,163 | 1.00 | 35 | 0.99998 |
| HiFi-MAG-Pipeline_complete.407 | ProxiMeta_bin_448 | 2,051,626 | 2,053,800 | 1.00 | 2,092,519 | 0.98 | 200 | 0.99990 |
| HiFi-MAG-Pipeline_complete.129 | HiFi-MAG-Pipeline_complete.698 | 2,492,326 | 2,493,722 | 1.00 | 2,483,383 | 1.00 | 24 | 0.99999 |
| ProxiMeta_bin_325 | ProxiMeta_bin_375 | 2,288,660 | 2,345,536 | 0.98 | 2,409,861 | 0.95 | 1156 | 0.99949 |
| HiFi-MAG-Pipeline_complete.578 | HiFi-MAG-Pipeline_complete.31 | 1,802,512 | 1,802,683 | 1.00 | 1,801,451 | 1.00 | 50 | 0.99997 |
| HiFi-MAG-Pipeline_metabat2.392 | ProxiMeta_bin_410 | 2,221,783 | 2,409,539 | 0.92 | 2,263,928 | 0.98 | 582 | 0.99974 |
| ProxiMeta_bin_451 | ProxiMeta_bin_461 | 1,850,061 | 1,917,498 | 0.96 | 2,046,665 | 0.90 | 1149 | 0.99938 |
| HiFi-MAG-Pipeline_complete.171 | HiFi-MAG-Pipeline_complete.690 | 1,713,777 | 1,706,528 | 1.00 | 1,706,314 | 1.00 | 21 | 0.99999 |
| HiFi-MAG-Pipeline_complete.136 | HiFi-MAG-Pipeline_metabat2.1363 | 2,812,401 | 2,839,231 | 0.99 | 2,812,453 | 1.00 | 61 | 0.99998 |
| HiFi-MAG-Pipeline_complete.308 | HiFi-MAG-Pipeline_semibin2_70 | 2,812,147 | 2,814,490 | 1.00 | 2,812,237 | 1.00 | 101 | 0.99996 |
| HiFi-MAG-Pipeline_complete.76 | ProxiMeta_bin_69 | 3,746,418 | 3,707,377 | 1.01 | 3,769,252 | 0.99 | 1087 | 0.99971 |
| HiFi-MAG-Pipeline_complete.322 | ProxiMeta_bin_283 | 2,765,499 | 2,954,804 | 0.94 | 2,765,564 | 1.00 | 60 | 0.99998 |
| HiFi-MAG-Pipeline_complete.78 | ProxiMeta_bin_210 | 3,038,040 | 3,019,444 | 1.01 | 3,049,825 | 1.00 | 44 | 0.99999 |
| HiFi-MAG-Pipeline_semibin2_6 | HiFi-MAG-Pipeline_semibin2_88 | 3,136,200 | 3,168,951 | 0.99 | 3,185,247 | 0.98 | 503 | 0.99984 |
| HiFi-MAG-Pipeline_complete.314 | HiFi-MAG-Pipeline_complete.68 | 1,425,786 | 1,409,852 | 1.01 | 1,410,124 | 1.01 | 1806 | 0.99873 |
| **HiFi-MAG-Pipeline_complete.343** | **HiFi-MAG-Pipeline_complete.584** | **1,631,970** | **1,631,756** | **1.00** | **1,631,967** | **1.00** | **7** | **1.00000** |
| **HiFi-MAG-Pipeline_complete.176** | **HiFi-MAG-Pipeline_complete.340** | **1,423,277** | **1,423,281** | **1.00** | **1,423,277** | **1.00** | **2** | **1.00000** |
| HiFi-MAG-Pipeline_metabat2.584 | HiFi-MAG-Pipeline_metabat2.1473 | 1,997,534 | 1,984,086 | 1.01 | 2,044,370 | 0.98 | 192 | 0.99990 |
| **HiFi-MAG-Pipeline_complete.418** | **HiFi-MAG-Pipeline_complete.919** | **3,252,302** | **3,252,301** | **1.00** | **3,252,299** | **1.00** | **5** | **1.00000** |
| ProxiMeta_bin_497 | ProxiMeta_bin_487 | 1,698,068 | 1,759,662 | 0.96 | 1,883,383 | 0.90 | 2014 | 0.99881 |
| HiFi-MAG-Pipeline_complete.388 | ProxiMeta_bin_195 | 2,948,729 | 2,968,664 | 0.99 | 3,117,025 | 0.95 | 622 | 0.99979 |
| ProxiMeta_bin_222 | ProxiMeta_bin_304 | 2,684,120 | 2,715,785 | 0.99 | 2,687,068 | 1.00 | 298 | 0.99989 |
| HiFi-MAG-Pipeline_complete.241 | HiFi-MAG-Pipeline_complete.446 | 2,380,652 | 2,380,546 | 1.00 | 2,381,043 | 1.00 | 63 | 0.99997 |
| **HiFi-MAG-Pipeline_complete.741** | **HiFi-MAG-Pipeline_complete.427** | **2,300,123** | **2,300,136** | **1.00** | **2,300,120** | **1.00** | **6** | **1.00000** |

| | | | | | | | | |
|---|---|---|---|---|---|---|---|---|
| HiFi-MAG-Pipeline_complete.354 | HiFi-MAG-Pipeline_complete.704 | 2,982,480 | 3,030,320 | 0.98 | 2,991,925 | 1.00 | 193 | 0.99994 |
| HiFi-MAG-Pipeline_complete.238 | HiFi-MAG-Pipeline_complete.174 | 2,610,790 | 2,624,232 | 0.99 | 2,665,448 | 0.98 | 363 | 0.99986 |
| ProxiMeta_bin_185 | ProxiMeta_bin_239 | 2,876,787 | 2,879,984 | 1.00 | 2,942,054 | 0.98 | 32 | 0.99999 |
| ProxiMeta_bin_170 | HiFi-MAG-Pipeline_complete.543 | 2,887,245 | 2,924,930 | 0.99 | 2,905,190 | 0.99 | 913 | 0.99968 |
| ProxiMeta_bin_554 | ProxiMeta_bin_549 | 1,522,674 | 1,571,980 | 0.97 | 1,528,212 | 1.00 | 36 | 0.99998 |
| HiFi-MAG-Pipeline_complete.276 | HiFi-MAG-Pipeline_complete.713 | 1,569,006 | 1,575,738 | 1.00 | 1,567,114 | 1.00 | 123 | 0.99992 |
| HiFi-MAG-Pipeline_complete.629 | HiFi-MAG-Pipeline_complete.301 | 2,364,693 | 2,364,636 | 1.00 | 2,365,167 | 1.00 | 85 | 0.99996 |
| HiFi-MAG-Pipeline_metabat2.575 | HiFi-MAG-Pipeline_complete.99 | 2,541,554 | 2,541,603 | 1.00 | 2,538,504 | 1.00 | 23 | 0.99999 |
| ProxiMeta_bin_214 | ProxiMeta_bin_262 | 2,703,034 | 2,752,519 | 0.98 | 2,853,051 | 0.95 | 263 | 0.99990 |
| ProxiMeta_bin_114 | ProxiMeta_bin_237 | 2,970,772 | 3,160,196 | 0.94 | 2,950,764 | 1.01 | 510 | 0.99983 |
| ProxiMeta_bin_34 | ProxiMeta_bin_40 | 4,005,449 | 4,077,766 | 0.98 | 4,216,776 | 0.95 | 58 | 0.99999 |
| HiFi-MAG-Pipeline_semibin2_48 | HiFi-MAG-Pipeline_semibin2_60 | 2,901,451 | 2,931,735 | 0.99 | 3,191,899 | 0.91 | 1105 | 0.99962 |
| **HiFi-MAG-Pipeline_complete.413** | **HiFi-MAG-Pipeline_complete.255** | **2,815,574** | **2,815,567** | **1.00** | **2,815,577** | **1.00** | **9** | **1.00000** |
| ProxiMeta_bin_135 | HiFi-MAG-Pipeline_complete.914 | 3,006,826 | 3,047,408 | 0.99 | 2,954,903 | 1.02 | 135 | 0.99996 |
| HiFi-MAG-Pipeline_complete.394 | HiFi-MAG-Pipeline_complete.452 | 2,064,646 | 2,068,472 | 1.00 | 2,065,531 | 1.00 | 960 | 0.99954 |
| HiFi-MAG-Pipeline_semibin2_87 | HiFi-MAG-Pipeline_semibin2_212 | 2,180,167 | 2,401,952 | 0.91 | 2,341,517 | 0.93 | 1207 | 0.99945 |
| HiFi-MAG-Pipeline_semibin2_178 | HiFi-MAG-Pipeline_semibin2_272 | 1,441,579 | 1,537,450 | 0.94 | 1,586,739 | 0.91 | 1366 | 0.99905 |
| **HiFi-MAG-Pipeline_complete.687** | **HiFi-MAG-Pipeline_complete.761** | **1,997,642** | **1,997,610** | **1.00** | **1,997,598** | **1.00** | **6** | **1.00000** |
| HiFi-MAG-Pipeline_metabat2.911 | HiFi-MAG-Pipeline_metabat2.308 | 3,114,290 | 3,140,354 | 0.99 | 3,114,942 | 1.00 | 202 | 0.99994 |
| HiFi-MAG-Pipeline_complete.745 | HiFi-MAG-Pipeline_complete.906 | 2,471,890 | 2,453,682 | 1.01 | 2,512,447 | 0.98 | 113 | 0.99995 |
| **HiFi-MAG-Pipeline_complete.389** | **HiFi-MAG-Pipeline_complete.353** | **1,623,728** | **1,623,738** | **1.00** | **1,623,825** | **1.00** | **101** | **0.99994** |
| HiFi-MAG-Pipeline_complete.645 | HiFi-MAG-Pipeline_complete.357 | 2,606,800 | 2,705,583 | 0.96 | 2,606,814 | 1.00 | 114 | 0.99996 |
| HiFi-MAG-Pipeline_complete.154 | HiFi-MAG-Pipeline_complete.927 | 3,772,415 | 3,772,303 | 1.00 | 3,772,490 | 1.00 | 33 | 0.99999 |
| ProxiMeta_bin_203 | ProxiMeta_bin_291 | 2,636,244 | 2,797,609 | 0.94 | 2,736,067 | 0.96 | 2708 | 0.99897 |
| **HiFi-MAG-Pipeline_complete.236** | **HiFi-MAG-Pipeline_complete.904** | **2,168,677** | **2,148,534** | **1.01** | **2,148,513** | **1.01** | **18** | **0.99999** |
| HiFi-MAG-Pipeline_complete.567 | HiFi-MAG-Pipeline_complete.910 | 2,481,173 | 2,488,750 | 1.00 | 2,450,412 | 1.01 | 11 | 1.00000 |
| **HiFi-MAG-Pipeline_complete.774** | **HiFi-MAG-Pipeline_complete.905** | **2,315,800** | **2,315,853** | **1.00** | **2,315,856** | **1.00** | **37** | **0.99998** |
| HiFi-MAG-Pipeline_complete.904 | HiFi-MAG-Pipeline_complete.913 | 2,808,882 | 2,809,810 | 1.00 | 2,808,288 | 1.00 | 77 | 0.99997 |

| | | | | | | | | |
|---|---|---|---|---|---|---|---|---|
| HiFi-MAG-Pipeline_complete.117 | ProxiMeta_bin_26 | 4,699,210 | 4,689,260 | 1.00 | 4,731,233 | 0.99 | 30 | 0.99999 |
| ProxiMeta_bin_183 | ProxiMeta_bin_292 | 2,696,748 | 2,882,291 | 0.94 | 2,733,624 | 0.99 | 1685 | 0.99938 |
| HiFi-MAG-Pipeline_metabat2.299 | HiFi-MAG-Pipeline_complete.606 | 3,724,797 | 3,704,825 | 1.01 | 3,678,284 | 1.01 | 76 | 0.99998 |
| HiFi-MAG-Pipeline_semibin2_26 | ProxiMeta_bin_137 | 3,107,395 | 3,236,376 | 0.96 | 3,362,455 | 0.92 | 4087 | 0.99868 |
| ProxiMeta_bin_173 | ProxiMeta_bin_190 | 2,828,819 | 2,916,014 | 0.97 | 3,136,612 | 0.90 | 2373 | 0.99916 |
| ProxiMeta_bin_120 | ProxiMeta_bin_218 | 2,951,556 | 3,131,901 | 0.94 | 3,015,585 | 0.98 | 3346 | 0.99887 |
| HiFi-MAG-Pipeline_metabat2.758 | HiFi-MAG-Pipeline_semibin2_57 | 2,689,311 | 2,688,221 | 1.00 | 2,756,686 | 0.98 | 172 | 0.99994 |
| ProxiMeta_bin_192 | ProxiMeta_bin_295 | 2,687,272 | 2,845,487 | 0.94 | 2,728,790 | 0.98 | 2066 | 0.99923 |
| HiFi-MAG-Pipeline_metabat2.932 | HiFi-MAG-Pipeline_metabat2.905 | 3,283,322 | 3,316,636 | 0.99 | 3,313,635 | 0.99 | 507 | 0.99985 |
| ProxiMeta_bin_85 | HiFi-MAG-Pipeline_complete.935 | 3,208,232 | 3,380,180 | 0.95 | 3,344,019 | 0.96 | 11656 | 0.99637 |
| ProxiMeta_bin_22 | ProxiMeta_bin_32 | 4,316,860 | 4,346,656 | 0.99 | 4,407,615 | 0.98 | 1678 | 0.99961 |
| HiFi-MAG-Pipeline_semibin2_39 | HiFi-MAG-Pipeline_semibin2_115 | 2,472,651 | 2,472,660 | 1.00 | 2,472,652 | 1.00 | 47 | 0.99998 |
| HiFi-MAG-Pipeline_metabat2.849 | ProxiMeta_bin_177 | 3,231,117 | 3,447,254 | 0.94 | 3,187,552 | 1.01 | 456 | 0.99986 |
| ProxiMeta_bin_16 | HiFi-MAG-Pipeline_metabat2.1470 | 4,403,211 | 4,522,536 | 0.97 | 4,491,594 | 0.98 | 2373 | 0.99946 |
| ProxiMeta_bin_66 | ProxiMeta_bin_92 | 3,326,751 | 3,588,007 | 0.93 | 3,658,037 | 0.91 | 877 | 0.99974 |
| ProxiMeta_bin_154 | HiFi-MAG-Pipeline_complete.921 | 2,974,660 | 2,978,324 | 1.00 | 3,136,417 | 0.95 | 208 | 0.99993 |
| ProxiMeta_bin_72 | ProxiMeta_bin_117 | 3,376,692 | 3,478,396 | 0.97 | 3,449,435 | 0.98 | 757 | 0.99978 |
| ProxiMeta_bin_142 | HiFi-MAG-Pipeline_semibin2_73 | 2,912,435 | 3,026,000 | 0.96 | 3,206,866 | 0.91 | 7542 | 0.99741 |
| HiFi-MAG-Pipeline_complete.218 | HiFi-MAG-Pipeline_semibin2_18 | 3,278,077 | 3,361,848 | 0.98 | 3,333,893 | 0.98 | 1895 | 0.99942 |
| ProxiMeta_bin_131 | ProxiMeta_bin_211 | 2,777,313 | 3,058,388 | 0.91 | 3,044,992 | 0.91 | 1013 | 0.99964 |
| ProxiMeta_bin_105 | ProxiMeta_bin_132 | 3,240,857 | 3,258,013 | 0.99 | 3,381,954 | 0.96 | 84 | 0.99997 |
| HiFi-MAG-Pipeline_complete.143 | HiFi-MAG-Pipeline_semibin2_106 | 3,212,644 | 3,247,179 | 0.99 | 3,392,372 | 0.95 | 553 | 0.99983 |
| ProxiMeta_bin_53 | ProxiMeta_bin_58 | 3,727,541 | 3,762,509 | 0.99 | 3,939,849 | 0.95 | 20 | 0.99999 |
| ProxiMeta_bin_91 | ProxiMeta_bin_139 | 3,153,263 | 3,353,722 | 0.94 | 3,353,434 | 0.94 | 2221 | 0.99930 |
| ProxiMeta_bin_55 | HiFi-MAG-Pipeline_metabat2.1225 | 3,666,548 | 3,746,642 | 0.98 | 3,653,221 | 1.00 | 272 | 0.99993 |
| HiFi-MAG-Pipeline_semibin2_25 | HiFi-MAG-Pipeline_metabat2.799 | 4,071,948 | 4,068,431 | 1.00 | 4,062,906 | 1.00 | 515 | 0.99987 |
| ProxiMeta_bin_414 | HiFi-MAG-Pipeline_semibin2_123 | 1,917,323 | 2,046,818 | 0.94 | 1,993,365 | 0.96 | 4262 | 0.99778 |
| **HiFi-MAG-Pipeline_complete.1031** | **HiFi-MAG-Pipeline_complete.945** | **6,096,093** | **6,096,109** | **1.00** | **6,096,089** | **1.00** | **12** | **1.00000** |

| | | | | | | | | |
|---|---|---|---|---|---|---|---|---|
| **HiFi-MAG-Pipeline_complete.507** | **HiFi-MAG-Pipeline_complete.358** | **2,459,755** | **2,459,752** | **1.00** | **2,459,775** | **1.00** | **20** | **0.99999** |
| HiFi-MAG-Pipeline_metabat2.508 | HiFi-MAG-Pipeline_semibin2_1 | 2,291,853 | 2,292,871 | 1.00 | 2,483,420 | 0.92 | 283 | 0.99988 |
| **HiFi-MAG-Pipeline_complete.387** | **HiFi-MAG-Pipeline_complete.121** | **1,910,996** | **1,910,992** | **1.00** | **1,910,995** | **1.00** | **4** | **1.00000** |
| HiFi-MAG-Pipeline_complete.48 | ProxiMeta_bin_179 | 3,028,789 | 3,013,614 | 1.01 | 3,170,620 | 0.96 | 124 | 0.99996 |
| ProxiMeta_bin_193 | HiFi-MAG-Pipeline_complete.165 | 2,784,795 | 2,843,320 | 0.98 | 2,965,240 | 0.94 | 10645 | 0.99618 |
| HiFi-MAG-Pipeline_complete.242 | ProxiMeta_bin_208 | 3,004,432 | 2,986,251 | 1.01 | 3,062,839 | 0.98 | 118 | 0.99996 |
| HiFi-MAG-Pipeline_metabat2.326 | HiFi-MAG-Pipeline_complete.653 | 2,579,337 | 2,593,052 | 0.99 | 2,596,052 | 0.99 | 147 | 0.99994 |
| HiFi-MAG-Pipeline_complete.91 | ProxiMeta_bin_18 | 5,199,871 | 5,199,887 | 1.00 | 5,313,050 | 0.98 | 238 | 0.99995 |
| HiFi-MAG-Pipeline_complete.419 | HiFi-MAG-Pipeline_complete.907 | 3,044,006 | 3,045,848 | 1.00 | 3,040,165 | 1.00 | 35 | 0.99999 |
| **HiFi-MAG-Pipeline_complete.436** | **HiFi-MAG-Pipeline_complete.94** | **2,515,111** | **2,515,134** | **1.00** | **2,515,110** | **1.00** | **16** | **0.99999** |
| HiFi-MAG-Pipeline_complete.366 | HiFi-MAG-Pipeline_complete.278 | 2,219,743 | 2,140,313 | 1.04 | 2,130,044 | 1.04 | 158 | 0.99993 |
| ProxiMeta_bin_86 | ProxiMeta_bin_94 | 3,316,043 | 3,379,963 | 0.98 | 3,652,674 | 0.91 | 26 | 0.99999 |
| HiFi-MAG-Pipeline_metabat2.1107 | HiFi-MAG-Pipeline_semibin2_22 | 4,157,286 | 4,262,062 | 0.98 | 4,572,019 | 0.91 | 1246 | 0.99970 |
| ProxiMeta_bin_3 | ProxiMeta_bin_5 | 6,654,779 | 6,761,406 | 0.98 | 6,738,685 | 0.99 | 66 | 0.99999 |
| HiFi-MAG-Pipeline_semibin2_7 | ProxiMeta_bin_19 | 5,162,894 | 5,132,555 | 1.01 | 5,291,314 | 0.98 | 275 | 0.99995 |
| HiFi-MAG-Pipeline_complete.247 | ProxiMeta_bin_138 | 3,334,608 | 3,318,890 | 1.00 | 3,358,009 | 0.99 | 571 | 0.99983 |
| **HiFi-MAG-Pipeline_complete.974** | **HiFi-MAG-Pipeline_complete.939** | **3,574,670** | **3,554,633** | **1.01** | **3,554,536** | **1.01** | **50** | **0.99999** |
| HiFi-MAG-Pipeline_metabat2.111 | HiFi-MAG-Pipeline_complete.806 | 2,338,894 | 2,412,602 | 0.97 | 2,339,377 | 1.00 | 136 | 0.99994 |

Matches in bold (n=17) indicate pairs that are each single contig, within 200 bp total length, and displaying <101 similarity errors.

**Supplemental Table S11.** Predicted mobile element contig counts from ProxiMeta, geNomad, and VirSorter2. Mobile contigs associated with host contigs are noted in parentheses.

| Viral detection method | Assembly method | Proviruses | Viruses | Total viral | Plasmids | Integrated plasmids | Total plasmids |
|---|---|---|---|---|---|---|---|
| ProxiMeta | hifiasm-meta | 795 | 1,380 (123) | 2,175 | 164 (25) | 2,268 | 2,432 |
| | metaMDBG | 728 | 1,773 (333) | 2,501 | 129 (32) | 1,878 | 2,007 |
| geNomad | hifiasm-meta | 2,199 | 2,352 | 4,551 | 6,827 | NA | 6,827 |
| | metaMDBG | 2,005 | 4,674 | 6,679 | 11,983 | NA | 11,983 |
| VirSorter2 | hifiasm-meta | 1,749 | 2,515 | 4,264 | NA | NA | NA |
| | metaMDBG | 1,737 | 4,466 | 6,203 | NA | NA | NA |

**Supplemental Figure S1. (a)** HiFi read length distributions for the Sequel IIe (top row) and Revio (bottom row) systems. (**b)** Number of complete passes per HiFi read (e.g., number of subreads). (**c)** Predicted per-read QV scores. The Sequel IIe summary is based on 17.15 million HiFi reads and 120.8 Gbp total data, and the Revio dataset contains 17.56 million HiFi reads and 135.0 Gbp total data.

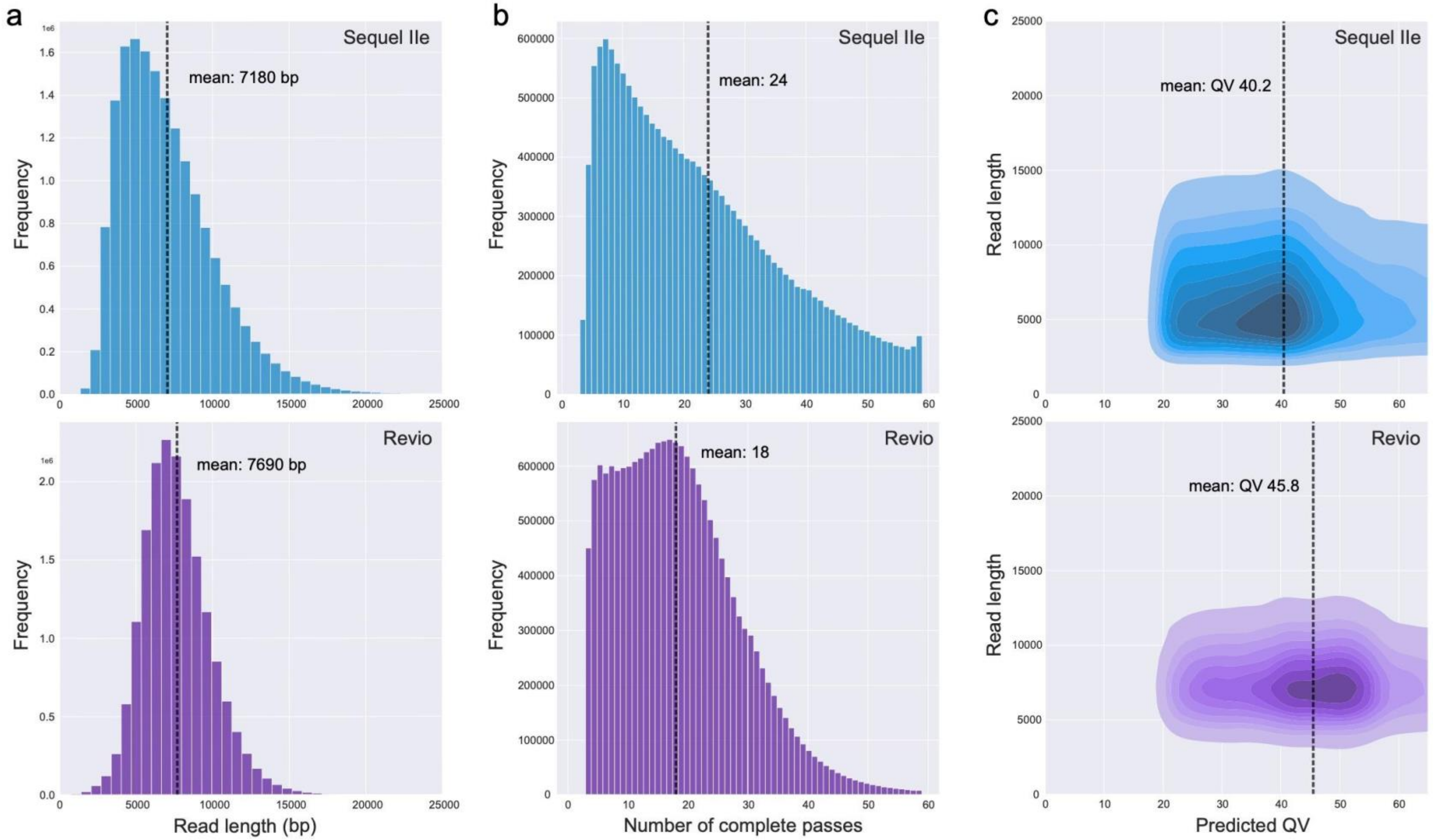

**Supplemental Figure S2.** Summary of quality characteristics for all categories of MAGs recovered by pb-MAG-mirror, across all combinations of assembly (hifiasm-meta, metaMDBG) and binning methods (HiFi-MAG-Pipeline, ProxiMeta).

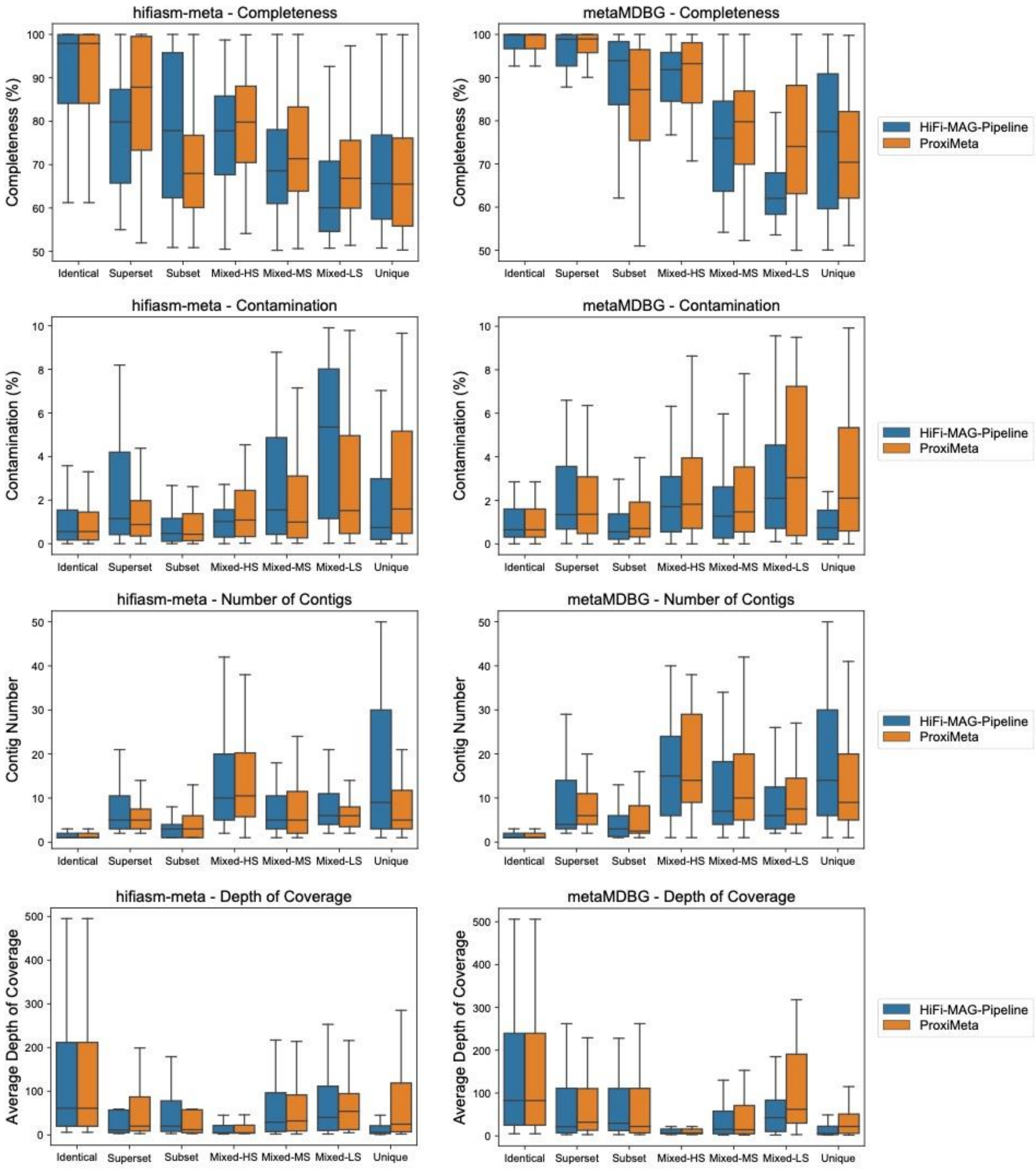

**Supplemental Figure S3.** Number of MAGs recovered for each downsampled dataset and binning method for hifiasm-meta. Total MAG numbers are shown above, with colors showing MAG counts exclusive to each category. HQ-MAGs require ≥90% single-copy genes (SCG) completeness and ≤5% contamination, whereas MQ-MAGs fall below the HQ thresholds but display ≥50% SCG completeness and ≤10% contamination.

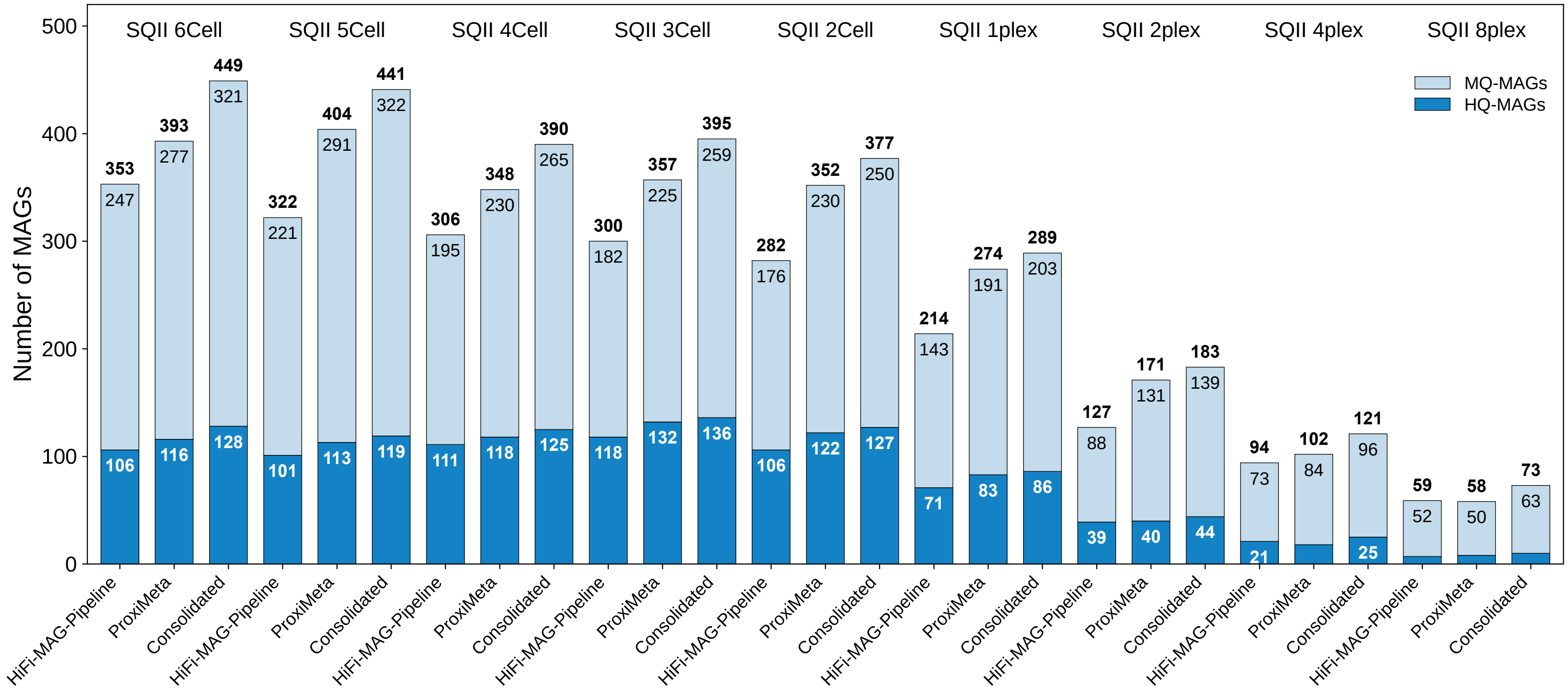

**Supplemental Figure S4.** Number of MAGs recovered for each downsampled dataset and binning method for metaMDBG. Total MAG numbers are shown above, with colors showing MAG counts exclusive to each category. HQ-MAGs require ≥90% single-copy genes (SCG) completeness and ≤5% contamination, whereas MQ-MAGs fall below the HQ thresholds but display ≥50% SCG completeness and ≤10% contamination.

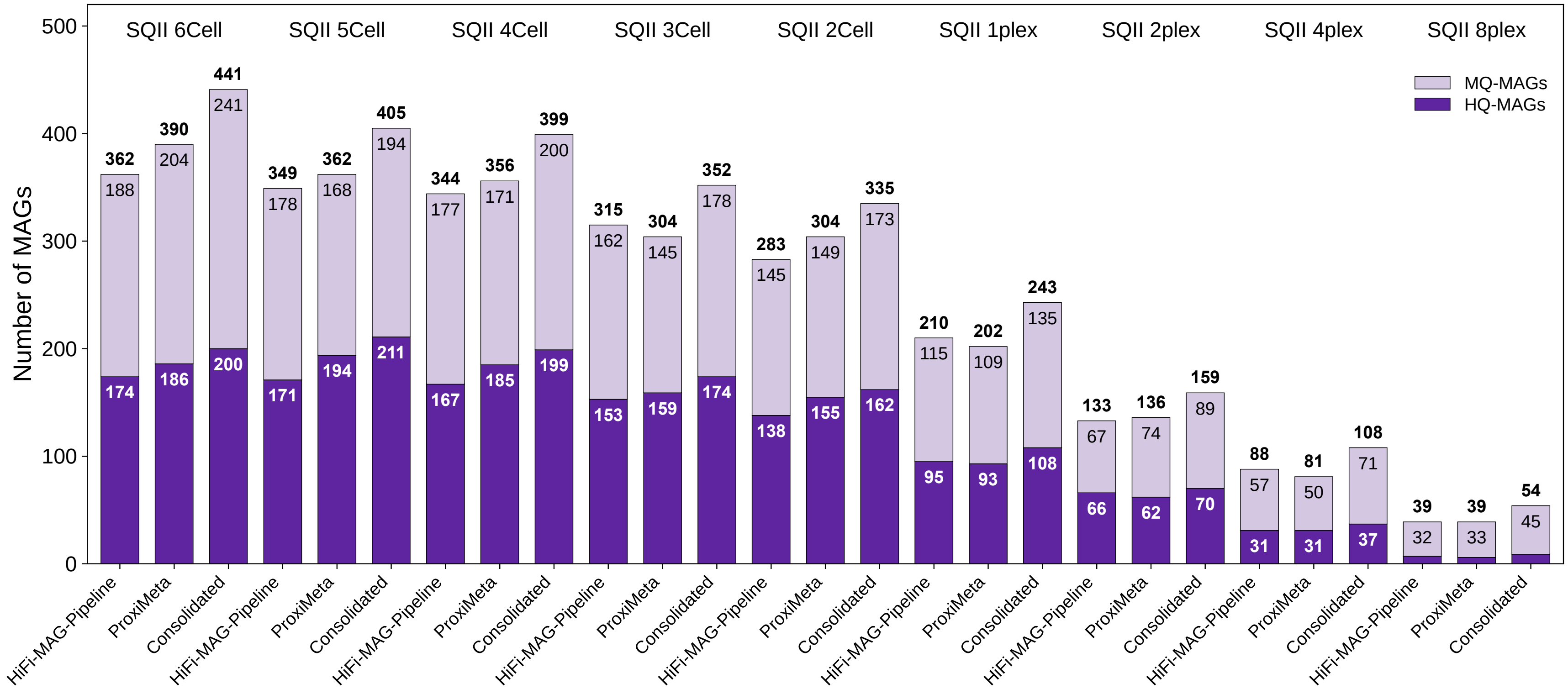

**Supplemental Figure S5.** Effects of minimum percent of genomes aligned and ANI on the number of unequivocally matched MAGs. Vertical line shows the percent genome aligned value (90%) used for reporting results.

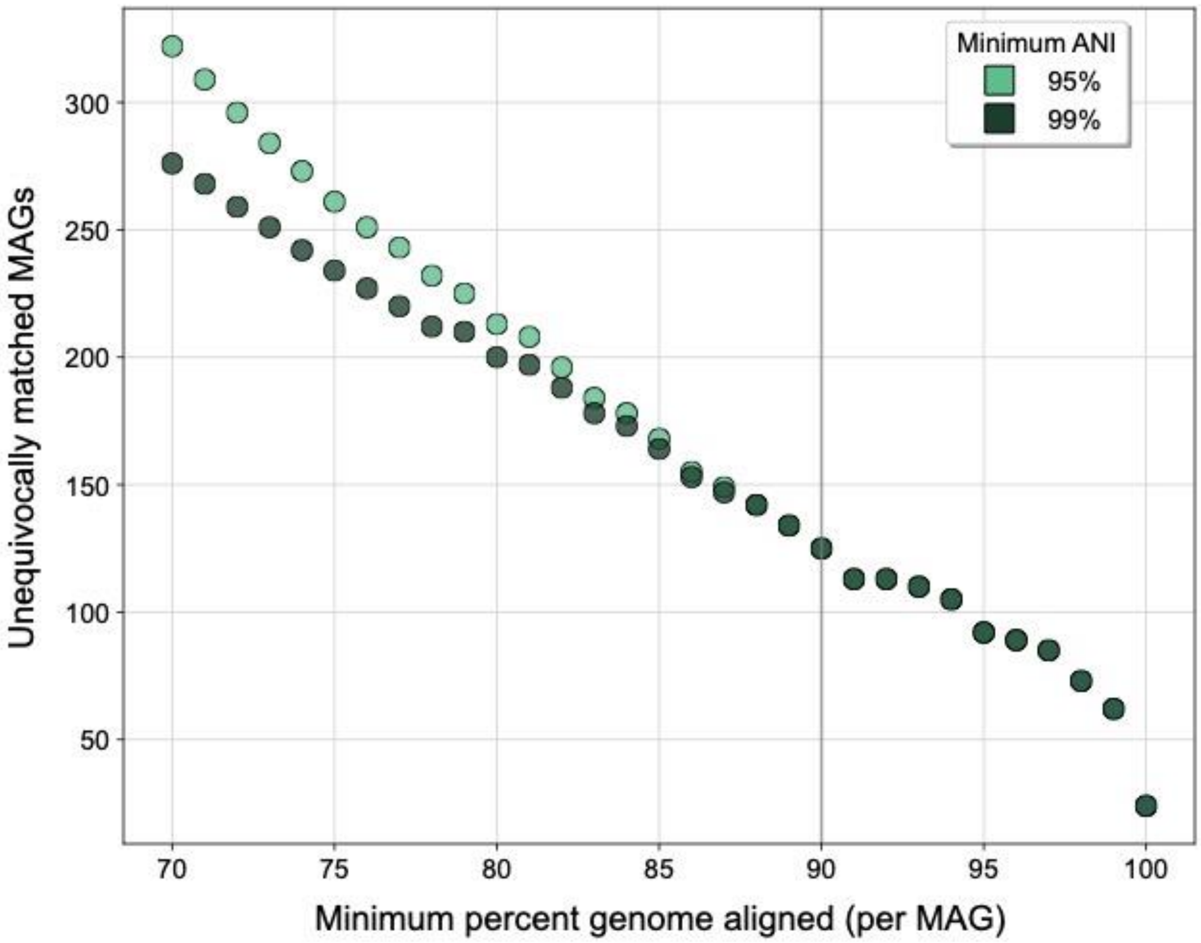

**Supplemental Figure S6.** Overlap of viral contigs across annotation methods for different classes.

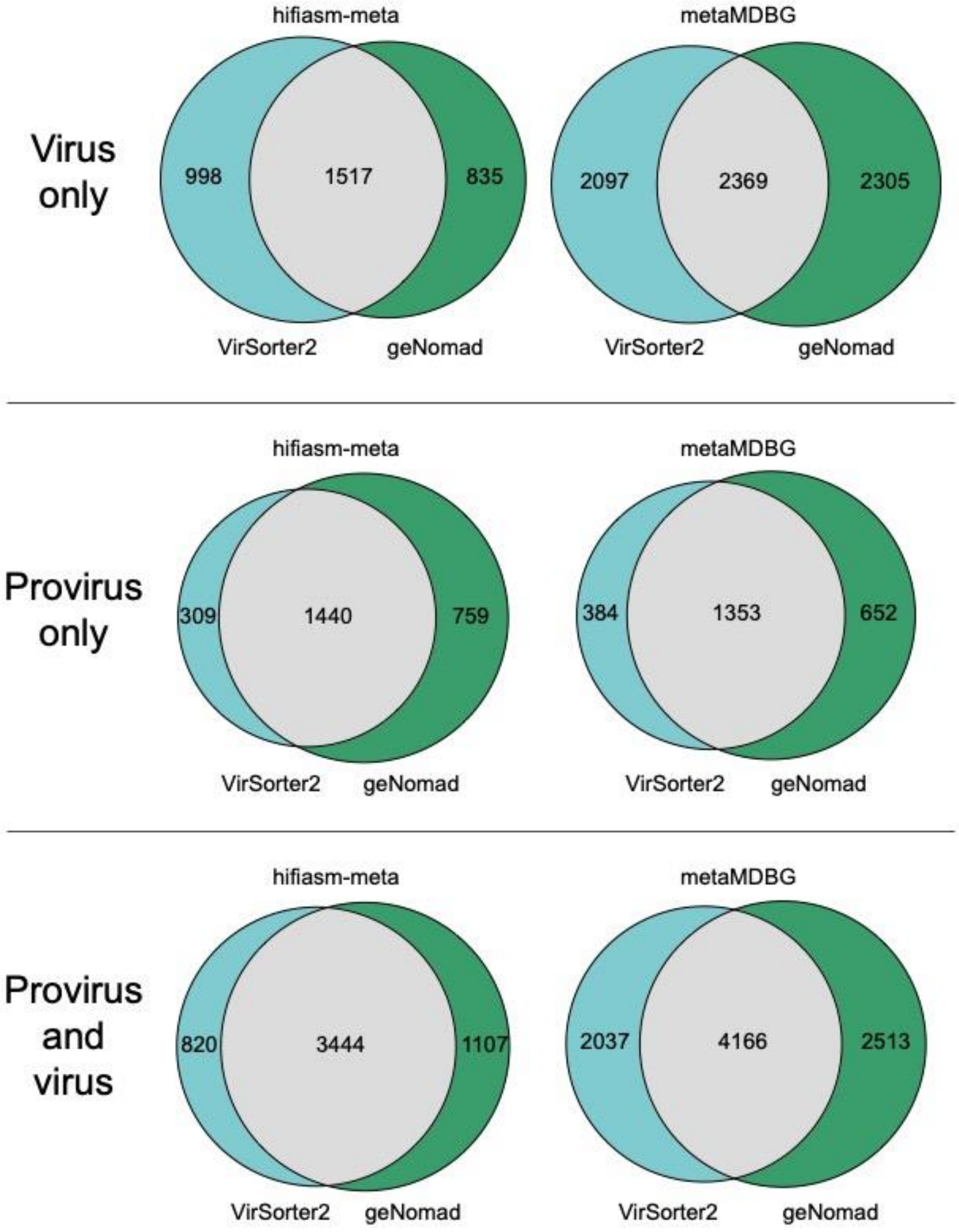
